# Omecamtiv mecarbil and Mavacamten target the same myosin pocket despite antagonistic effects in heart contraction

**DOI:** 10.1101/2023.11.15.567213

**Authors:** Daniel Auguin, Julien Robert-Paganin, Stéphane Réty, Carlos Kikuti, Amandine David, Gabriele Theumer, Arndt W. Schmidt, Hans-Joachim Knölker, Anne Houdusse

## Abstract

Inherited cardiomyopathies are amongst the most common cardiac diseases worldwide, leading in the late-stage to heart failure and death. The most promising treatments against these diseases are small-molecules directly modulating the force produced by β-cardiac myosin, the molecular motor driving heart contraction. Two of these molecules that produce antagonistic effects on cardiac contractility have completed clinical phase 3 trials: the activator *Omecamtiv mecarbil* and the inhibitor *Mavacamten*. In this work, we reveal by X-ray crystallography that both drugs target the same pocket and stabilize a pre-stroke structural state, with only few local differences. All atoms molecular dynamics simulations reveal how these molecules can have antagonistic impact on the allostery of the motor by comparing β-cardiac myosin in the apo form or bound to *Omecamtiv mecarbil* or *Mavacamten*. Altogether, our results provide the framework for rational drug development for the purpose of personalized medicine.

## Main text

Inherited cardiomyopathies are a major health concern, being one of the most important cause of heart disease worldwide. Heart failure due to end-stage cardiomyopathies can lead to sudden deaths^1,2,3^. These diseases are associated with single point mutations of contractile proteins from the sarcomere, such as β-cardiac myosin^4^. The effect of these point mutations has been extensively studied, but remains poorly understood. Up to now, therapeutic approaches to treat end-stage inherited cardiomyopathy have been highly invasive, including cardioverter-defibrillator implantations and heart transplantation^5^.

In the heart, β-cardiac myosin is the major nanomotor that produces force during contraction. Myosins are ATP-dependent molecular motors involved in almost all processes of life (reviewed by Houdusse et al.^6^). The force produced depends in particular on the regulation of β-cardiac myosin that controls the number of active nanomotors. The double-headed β-cardiac myosin can indeed adopt an inactive, sequestered state, unable to participate in force production, which corresponds to a specific motif called the interacted-heads motif (IHM)^7,8,9^. When exertion increases, destabilization of the sequestered state allows an increase in the number of myosins engaged in force production^10^.

Small-molecules directly targeting β-cardiac myosin can modulate force production and are promising approaches to treat cardiac diseases^11,12^. Sarcomere activators can increase contractile force^13^, while inhibitors of myosin force production can decrease the force produced when the heart is hypercontractile^14,15^. Two modulators have completed phase 3 clinical trials. The activator of sarcomere contraction *Omecamtiv mecarbil* (OM), led to a reduction in mortality and cardiovascular events in the treatment of heart failure with reduced ejection fraction (HFrEF), (GALACTIC-HF^13^). Another modulator, the inhibitor *Mavacamten* (Mava), became the first revolutionary FDA-approved treatment, and is now available for patients of obstructive hypertrophic cardiomyopathies (HCM) under the name of CAMZYOS^TM^ (FDA, 2022).

These two molecules have antagonistic effects on the heart: while OM is able to increase cardiac contractility without side effects on calcium concentrations or myocardial oxygen consumption^16,17^; Mava is able to decrease cardiac contractility and to suppress hypertrophy and cardiomyocyte disarray^14^.

The mechanism of action of OM includes the fact that it destabilizes the sequestered state, thus increasing the number of heads engaged in force production^18^. In contrast, Mava is reported to stabilize the folded-back sequestered state^19,20^ (**Fig. 1a, 1b**). Although direct measurements by FRET indicated that Mava increases the proportion of IHM by only 3% in an isolated doubled-headed myosin fragment (HMM)^21^. The contribution of these modulators on the motor cycle must also play an important role in their mechanism to modulate force contraction. OM increases the actin-activated ATPase activity and P_i_ release rate but also time spent on actin^22,23^. In contrast, Mava decreases the actin-activated ATPase activity, P_i_ release rate and affinity for actin^14,24^. We solved the X-ray structure of OM-bound β-cardiac myosin. This revealed the allosteric binding pocket of the drug and how it stabilizes the pre-powerstroke state (PPS), a state with a primed Lever arm, in which hydrolysis products are trapped until binding to F-actin triggers their release^25^. Importantly, recent single-molecule data showed that OM inhibits the working stroke of the wild-type (WT) β-cardiac myosin while it increases the working stroke of the HCM mutant R712L^26,27^.

**Figure 1.**
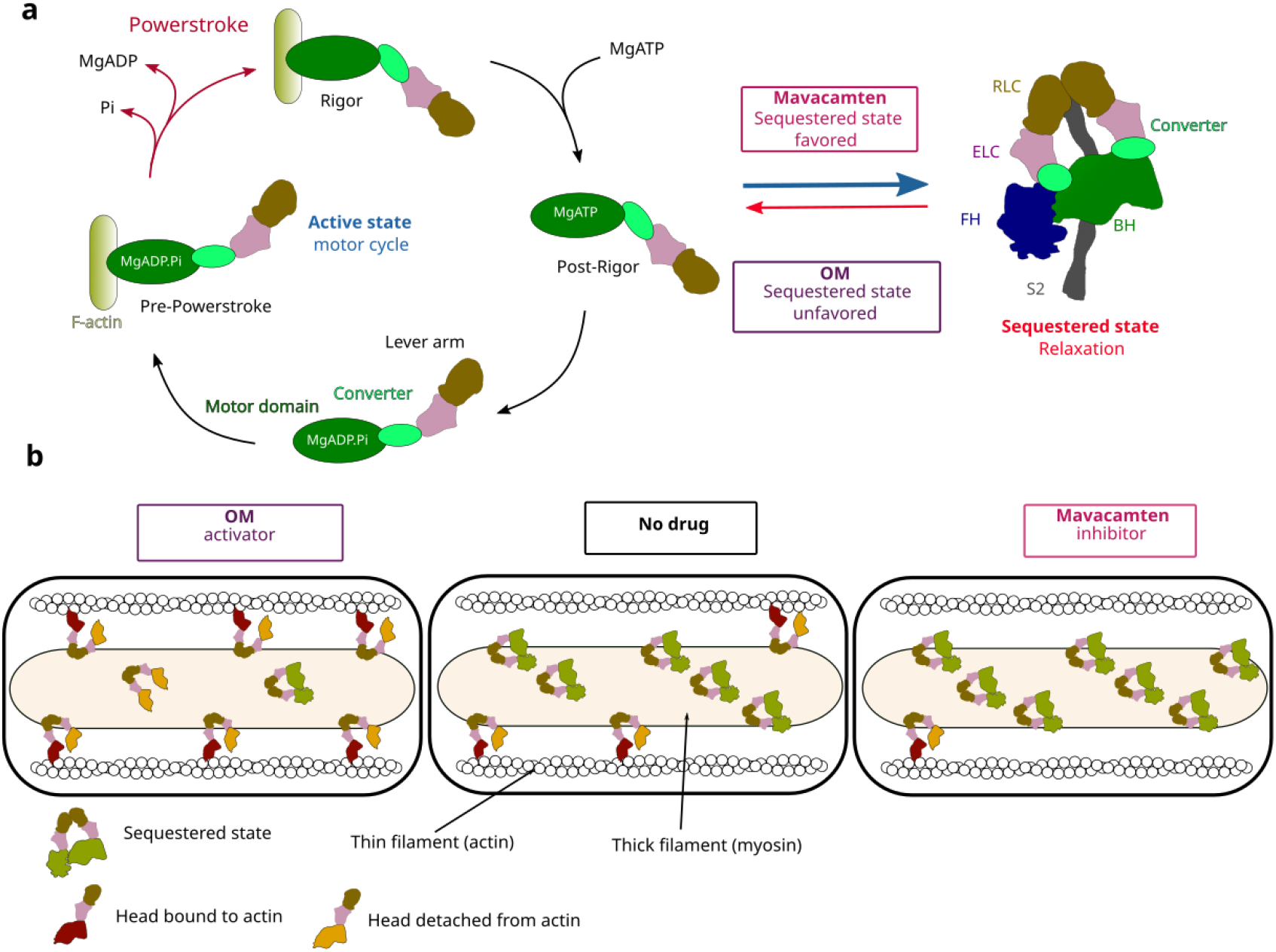
– Effect of Omecamtiv mecarbil (OM) and Mavacamten (Mava) on the motor cycle and regulation of β-cardiac myosin. **(a)** During its motor cycle, the myosin goes through three major structural states, associating to the actin in the pre-powerstroke (PPS) with the Lever arm up and performing the powerstroke which is responsible for heart contraction. In cardiac sarcomere, the population of available myosin is highly regulated by the presence of an inactive sequestered state that shuts down a proportion of myosin to regulate force production during contraction phases. OM and Mava have antagonistic effects on both the motor cycle and sequestered state stability. Mava is able to stabilize the sequestered state while OM destabilizes it, increasing the proportion of available heads. **(b)** On the active state, OM stabilizes the PPS state, increasing the number of heads available to interact with actin^25^, whereas Mava inhibits both the interaction with actin and the Pi release rate with an unknown mechanism.

Despite extensive efforts in characterizing these modulators, their precise mode of action remains unclear, including the rationale of how they can antagonistically influence the force output of the heart, and the actin-activated P_i_ release rate. This question could not be addressed so far since the binding site for Mava was unknown.

Structural information is also needed to design new modulators capable of enriching a personalized medicine approach. Indeed, hundreds of point-mutations are responsible for inherited cardiomyopathies and these mutations have diverse effects on β-cardiac myosin function and/or regulation^28,29,30,31^. Some mutations may impede the action of modulators, either by (i) reducing their affinity or (ii) blocking their allosteric effect. For example, the efficiency of OM is reduced in the case of the dilated cardiomyopathy (DCM)-causing mutation F764L^32^. It is thus mandatory to dissect the mode of action of OM and Mava on β-cardiac myosin, as this will guide the generation of novel drug candidates allowing all classes of mutations causing inherited cardiomyopathies to be treated.

In this work, we solved the structure of Mava-bound bovine β-cardiac myosin. We unexpectedly found that the allosteric binding pocket of this inhibitor is also where OM, the activator of contraction, binds. All-atom molecular dynamics simulations explains the source of their antagonistic effect on myosin allostery. This provides the blueprint of the mechanisms underlying modulation of myosin activity and regulation through the OM/Mava pocket.

## Results

### Mavacamten stabilizes a pre-stroke state

We successfully solved the structures of (i) the S1 fragment complexed with Mava and Mg.ADP.BeFx (PPS-S1-Mava), (ii) the proteolyzed motor domain fragment (MD) complexed with Mavacamten and Mg.ADP.BeFx (PPS-MD-Mava) and (iii) the structure of the motor domain fragment complexed to Mg.ADP.Vanadate (PPS-MD-Apo) at a resolution of 2.61; 1.80 and 2.76 Å respectively (**Supplementary Table 1&2**). Reprocessing of the *Omecamtiv mecarbil*-bound dataset^25^ with the StarAniso procedure^33^ provided a 1.96 Å resolution structure (PPS-S1-OM). The drugs were identified without ambiguity in the electron density map (**Supplementary Fig. 1**). Despite the different space groups and environments, the Mava-bound structures are nearly identical (Fig. 2a), which is a strong validation of the binding site. Like OM, Mava co-crystallizes with cardiac myosin in a pre-powerstroke (PPS) state, a state that traps the hydrolysis products with a primed Lever arm. Visualizing Mava and OM bound in the PPS state is relevant and consistent with the transient kinetic studies^25,24,23^, as these drugs are compatible with ATP hydrolysis, yet slow P_i_ release in the absence of actin reducing basal ATPase. No major difference is found in the PPS conformation upon drug binding, except for the position of the Converter (**Fig. 2a, Supplementary Fig. 2**).

**Figure 2.**
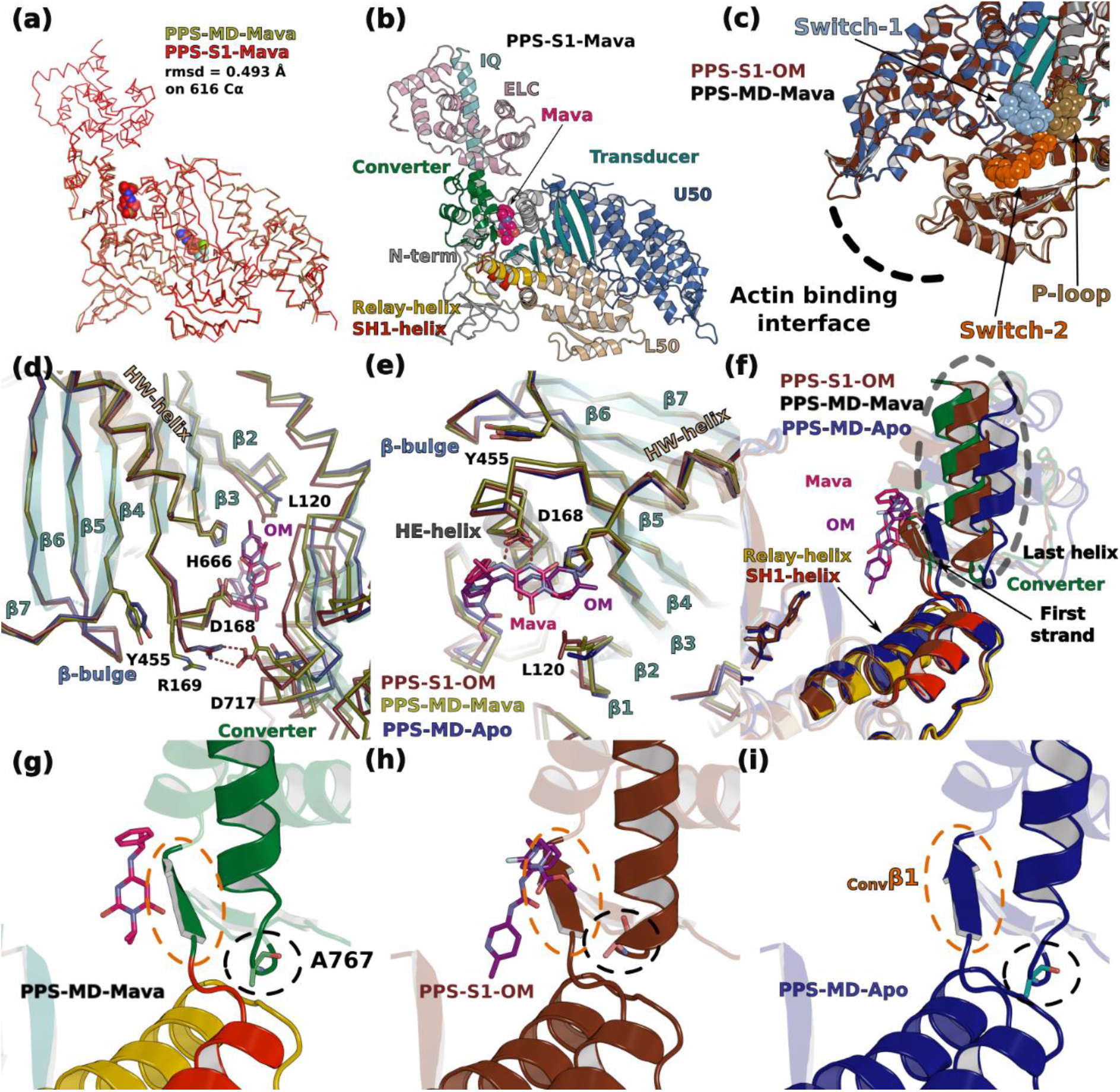
– Crystal structures of β-cardiac myosin complexed with Mavacamten and comparison with other PPS structures. **(a)** The motor domain (MD) and the S1 fragment (S1) structures complexed to Mavacamten (Mava) superimpose with a rmsd of 0.493 Å on 616 Cα. **(b)** Overall structure of the S1 fragment of β-cardiac myosin bound to Mava. The compound occupies a pocket between the Lever arm and the motor domain. **(c)** Superimposition of the OM bound and the Mava bound structures on the U50 shows that there is no difference of conformation on the actin binding interface and on the three nucleotide binding loops of the active site: Switches-1 and −2 and P-loop (spheres). **(d)**, **(e)** and **(f)** Local conformational differences induced by the presence of OM or Mava, notably in linkers near the Transducer and in the Lever arm orientation. The Lever arm is more primed in the structures in which either of these drugs are bound, compared to the apo structure. **(g)**, **(h**) and **(i)** Conformation of Ala767 and of the first β-strand of the Converter (_Conv_β1) depending on the presence of drugs.

### OM and Mava target the same pocket

Unexpectedly, both drugs target the same pocket, located between the N-terminal (N-term) and the Converter subdomains of the motor domain (Fig. 2b). No differences are found in the active site, including in the Switch-2 position (Fig. 2c). These structures are also similar for the conformation of the internal pocket – so called 50 kDa cleft – whose closure/opening controls the affinity for the actin filament (Fig. 2c). Such a similarity between the OM- and the Mava-bound structures is unexpected since OM and Mava strikingly differ in the way they increase (OM) or reduce (Mava) the rates of actin-activated P_i_ release^22,23,24^. A close comparison of the three PPS structures reveals that the differences are small and exist mainly around the drug binding site: **(i)** local differences in the ^Nter^HE-helix and the preceding linker (both involved in drug binding), (see D168 and R169, **Fig. 2d-e**), extend to small differences in the ^Transducer^β4 strand (see Y455 in **Fig.2d-e**), near the ^Transducer^HO-linker and the ^Transducer^beta-bulge **(ii)** non identical interactions of the drugs with the Converter (**Fig. 2d, 2f; Supplementary Movie 1**) also lead to local differences in particular for the position of ^Conv^Ala767 at the beginning of the last helix of the converter (**Fig. 2 g-i**). OM stabilizes the first turn of this helix but not Mava (**Supplementary Fig. 1**). In the two drug-bound structures, closure of the allosteric pocket around the drug results in a similar priming of the Lever arm angle for the two drugs (∼78° for OM and Mava, versus ∼69° for apo) (**Supplementary Fig. 3a-f; Supplementary Movie 1**). Differences in drug interactions with the Converter also translate to differences in the Relay, SH1-helix and Converter orientations (**Supplementary Fig. 4**), while distinct interactions of ^HW^H666 with both drugs lead to small effects on the first two Transducer strands (β1 and β2, near L120, **Fig. 2d**). The end of the ^Transducer^β2 strand (L120) uniquely participates in Mava binding. In addition, the size and conformation of the drug-binding pocket differ between OM, Mava and the apo structures (**Supplementary Fig. 3g-l**). Only in the most closed configuration found when OM binds, does the beginning of the β4 strand (R169) interact with ^Conv^D717 (**Fig. 2d**). Thus, important distinction in the drug interactions affect both the Transducer and the Lever arm, which are key elements of the allosteric communication in myosin motors, involved in force generation^6^.

### Interactions mediated by OM and Mava in the crystal structures

To compare how Mava and OM bind in this pocket, the drugs moieties were delineated and named A to C for Mava and A to D for OM (**Supplementary Fig. 5**). The drugs mediate similar polar bonds with critical residues of the two pockets (**Fig. 3a, 3b, Supplementary Table 3**): direct interactions involve ^N-term^Asp168, the amide nitrogen of the ^Conv^Arg712 backbone and the side chain of ^Conv^Asn711 with the carbamoylamino OM and the amino and pyrimidine groups of Mava. Mava and OM are also involved in similar apolar interactions (**Fig. 3a, 3b, Table 3**). Few additional residues differ between the OM and Mava pockets since OM is thinner but longer than Mava (**Fig. 3a, 3b**). The isobutyl group of Mava uniquely interacts with ^N-term^Leu120 (**Fig. 2d, 3a**). In contrast, the methyl ester piperazine ring of OM uniquely reaches residues of the linker prior to the HE helix as well as the last helix of the Converter on one end (K146, R147, ^HE^N160, ^Conv-H3^A767), while the methyl-pyridine reaches the Relay (H492, E497) on the other end of the OM molecule (**Fig. 3a, 3b**).

**Figure 3.**
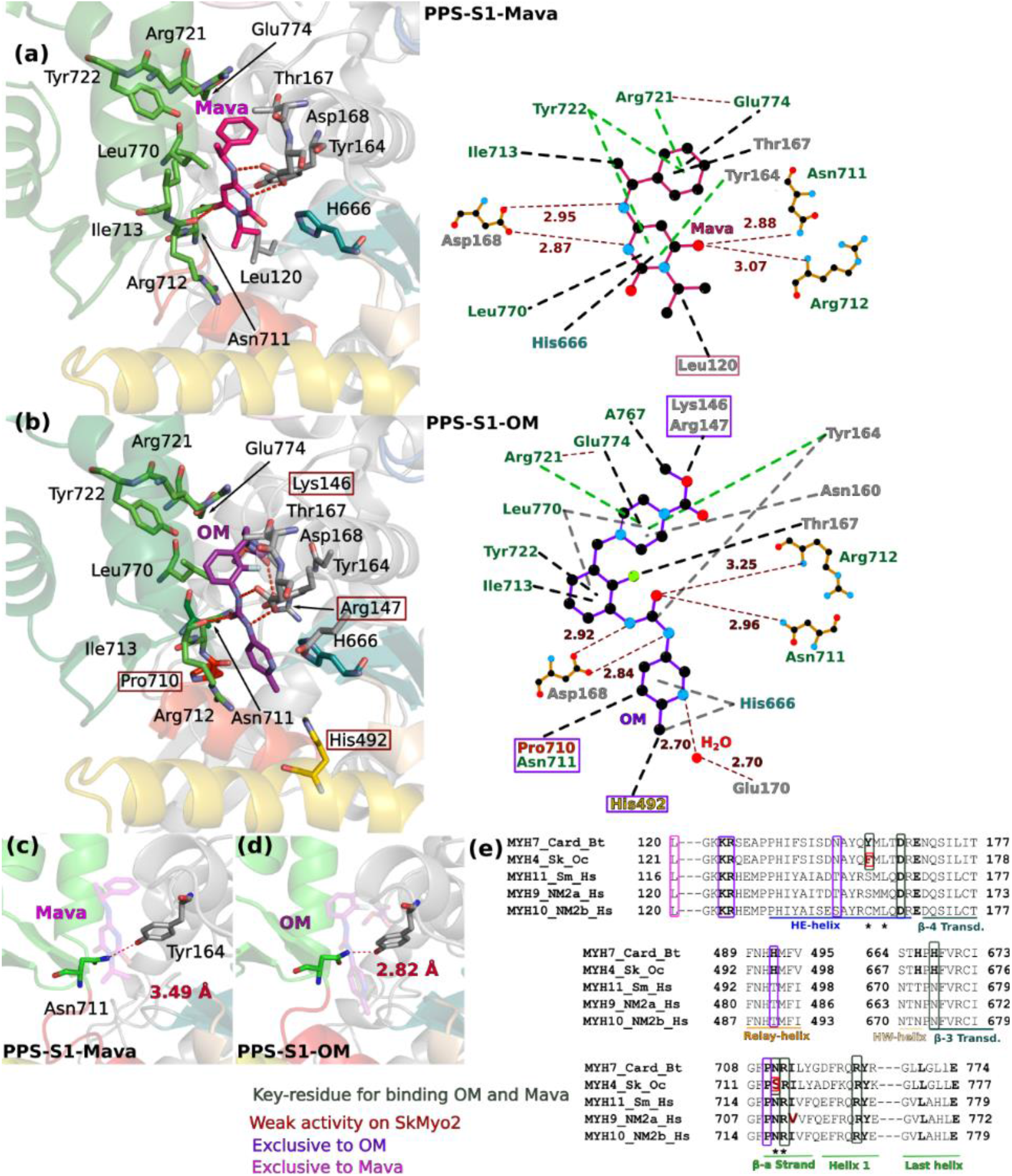
– Mava and OM targets the same pocket. **(a)** Cartoon and schematic representation of Mava binding site. **(b)** Cartoon and schematic LigPlot based^34^ representation of OM binding site. **(c)** and **(d)** Distinct shapes of the pocket where Mava and OM bind leads to different distances between the Converter and the motor domain, as shown by the distance between Y164 and N711 in the drug-bound structures. **(e)** Sequence alignment of class 2 myosins highlights the specificity determinants of OM and Mava. Residues important for binding both drugs are boxed in green, residues that lead to weaker affinity for skeletal myosin are colored in red, residues exclusive to OM or Mava binding are colored in deep purple and pink respectively. A star below the sequence indicates residues involved in electrostatic interactions

Sequence differences among different myosin-2s for residues around this pocket correlate with drug specificity. As OM, Mava is specific for α- and β-cardiac myosin. Fast skeletal myosin is inhibited with a ten-fold higher IC_50_, while smooth and non-muscle myosin-2 are not inhibited^24^. The weaker activity for fast skeletal myosin comes from only two small sequence differences in residues that directly bind the drugs (^Card^Y164/^Sk^F, ^Card^N711/^Sk^S) (**Fig. 3e; Supplementary Fig.6**). In contrast, several sequence differences are responsible for the inability of Mava or OM to target other myosin-2 (Fig. 3e).

### Mobility of the primed Lever arm greatly differs when Mava or OM are bound

To decipher how small local differences near the binding site of these force modulators can translate into different energy landscapes and thus different control of actin-binding or P_i_ release from the motor, all atoms molecular dynamics simulations were performed. Such simulations provide indications for allosteric communication and subdomain reorganization when conformational sub-states are explored^30,35,36,37,38,39,40^. We made a comparative study between three simulations started from S1 bound to Mava, S1 bound to OM and S1 without compound (apo), all bound to Mg.ADP.P_i_ in the active site (**Supplementary Movie 2, 3 and 4**). To guarantee a reproducibility of the results and minimize artifacts, the simulations were performed at least twice, from independent minimizations of the structures (See Material and Methods).

We first monitored the Lever arm position by plotting the root-mean-square displacement (RMSD) for backbone Cα atoms relative to their initial minimized complex structures during each simulation (Cα RMSD plot, Fig. 4a). In the Apo condition, the Lever arm swings back and forth exploring conformations differing by up to ∼36° (**Fig. 4a, Supplementary Fig. 7, 8 and Supplementary Movie 2**). When OM and Mava are bound, the amplitude between the extreme positions explored by the Lever arm are 13° and 26° respectively (**Fig. 4b, 4c**). The Lever arm is the most mobile in the Apo condition, with large movements leading to explore less primed Lever arm states. Mava slightly restrains the amplitude of the movements (**Supplementary Fig. 8, 9**). In contrast, OM binding greatly restrains the Lever arm movements and maintains it highly primed (**Supplementary Fig. 8, 9**). This is consistent with SAXS studies^25^ and single molecule experiments^26^ (that described that OM maintains the Lever arm up. The presence of OM and Mava both exert a cohesive action between the Lever arm and the motor domain that contributes to maintain the Lever arm primed (**Supplementary Movie 2, 3 and 4**). However, OM and Mava differ greatly in their mobility within the pocket: OM is kept for longer time in a specific position (close to that found in the crystal structure, **Supplementary Fig. 10, Supplementary Movie 5**), while Mava explores more easily different positions in the pocket (**Supplementary Fig. 9, Supplementary Movie 6**). Distinct fluctuations when Mava or OM are bound in the pocket correspond to distinct possible movements of the Converter/Lever arm orientation and positions explored (**Supplementary Movie 7**).

**Figure 4.**
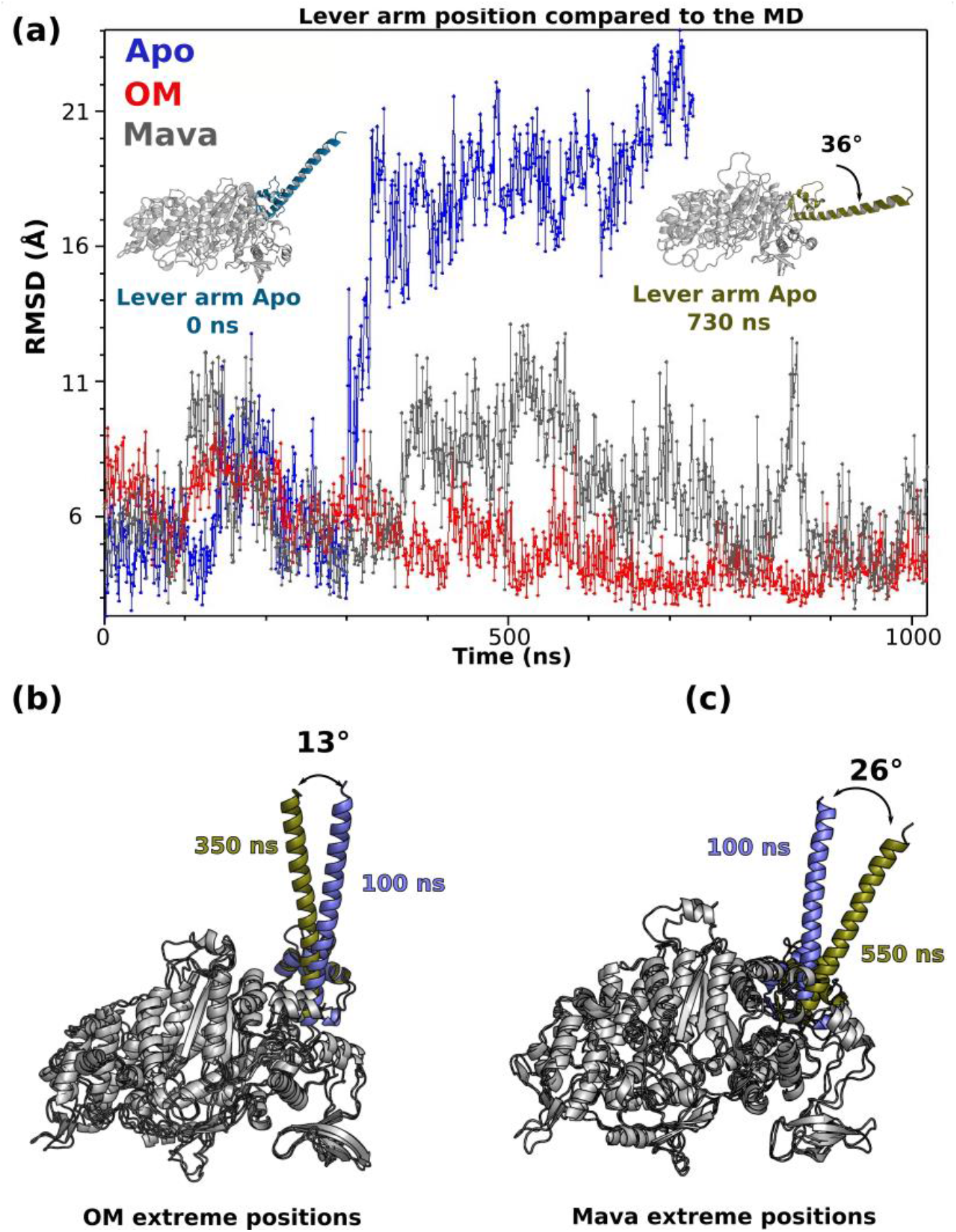
– OM and Mava stabilize the priming of the lever arm. **(a)** Evolution of the root mean square deviation (RMSD) of the Cα positions of the Lever arm (711-806) compared to the motor domain (MD) (1-710) for the three conditions during the time course of the simulation: Apo (blue), OM (red) and Mava (stone grey). The two extreme positions of the Apo (0 ns and 730 ns) are represented in blue and sand green, showing that a 36° swing occurred during the simulation. **(b)** Cartoon representation of the OM and Mava conditions which displays the extreme positions of the Lever arm. Mava allows more flexibility in Lever arm condition than OM. All the structures were aligned on the N-term subdomain for these measurements

### Distinct effects of Mava and OM on motor domain allostery

Since the actin-dependent P_i_ release rate is activated by OM and decreased by Mava, we then analyzed the dynamics within the motor domain, and in particular how the relative position of nucleotide-binding elements (P-loop, Switch-1 and Switch-2) behave along the simulations. These elements constitute the “backdoor”, which is closed in the pre-powerstroke state, but that must open during actin-activated P_i_ release^41,6^. The dynamics within the motor domain is of lower amplitude compared to that found for the Lever arm, as shown by the Cα RMSD plot (Fig. 5a). Interestingly, while the Apo and OM conditions follow similar dynamics, Mava is the most agitated and its dynamics greatly differs (Fig. 5a). In order to monitor the stability of the actin-binding interface that involves elements from the U50 and the L50, we followed the relative dynamics of the Helix-Turn-Helix (HTH) (Fig. 5b) and of the HO-helix in each condition (Fig. 5c). The HTH is part of the actin-binding interface and an element of the L50 and the position fluctuation of the long HO-helix of the U50 can be a reporter of the dynamics and position of this subdomain.

**Figure 5.**
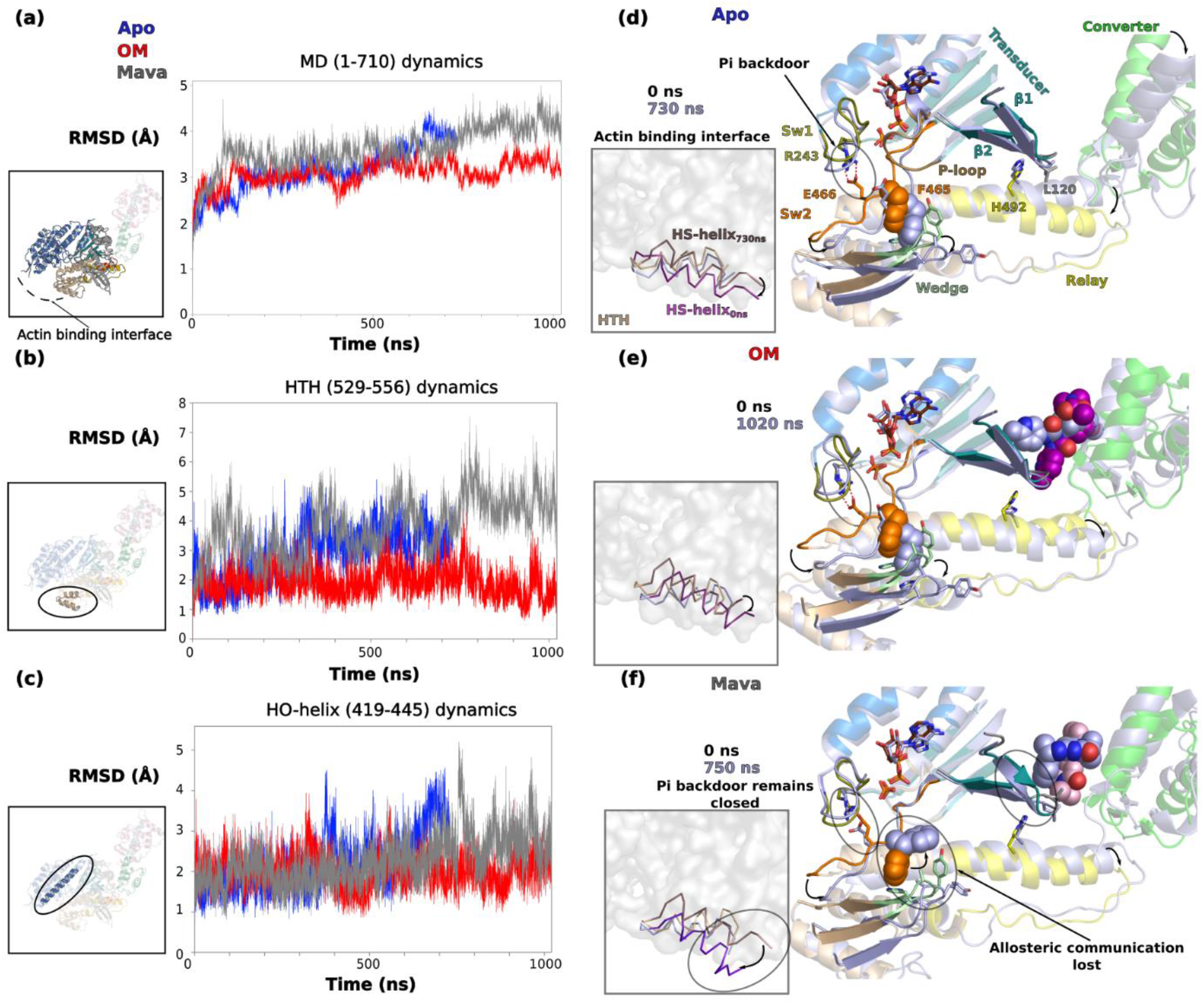
– OM and Mava induce different effects on the allostery of the motor. **(a)** RMSD of the Cα positions of the motor domain (1-710) during the time course of the simulation for the three conditions: Apo (blue), OM (red) and Mava (grey). Following the same methodology, **(b)** and **(c)** RMSD of the Cα positions of the helix-turn-helix (HTH, 529-526) and of the HO-helix (419-445), respectively. In **(d)**, **(e)** and **(f)**, the extreme positions of the HTH (part of the actin binding interface, the HS-helix is colored differently) is represented as well as that of the connectors for Apo, OM and Mava, respectively. The P_i_ backdoor is found close to the salt bridge between Switch-1 R243 and Switch-2 E466. In both the Apo and the OM conditions, the communication between the Wedge and the Switch-2 remains during all the duration of the simulation through the interaction with _Sw2_F465. This communication stabilizes the conformation of the HTH and of the actin binding interface. In the Mava condition, the communication between the Wedge and Switch-2 is rapidly lost in the simulation by the flip of F465, leading to a drastic change in the conformation of the HTH and specifically the HS-helix **(f)**. When Mava is bound, the P_i_ backdoor remains closed during all the simulation.

The dynamics of the connectors in the apo condition reveals that opening of the backdoor is explored via a Switch-2 movement that leads to the loss of the salt-bridge between Switch-1 and Switch-2 (R243/E466) (**Fig. 5b, Supplementary Fig. 7, Movie 2**). These fluctuations represent the first exploration of a transition towards the P_i_ release state, without much change in the L50 subdomain conformation and only small changes for the outer internal pocket (**Fig. 5b, Movie 2**). In the OM simulation, the backdoor displays high dynamics but does not fully open, as it is the case in the apo condition (**Fig. 5e, Supplementary Fig. 9a-c, Movie 2, 3**). For apo and OM, the communication between Switch-2 and the ^L50^Wedge is maintained via F465 (**Fig. 5b-c**), which restrains the L50 subdomain orientations in a PPS-like position. The overall landscape of states explored during the simulations of both apo and OM describe small-range fluctuations at the actin interface that are coupled with a dynamic back door that keeps P_i_ trapped. These data are consistent with an exploration of states along the OM simulation that are similar to those of apo, as they do not deviate much from the initial PPS conformation, with minimal changes in the actin interface since the position of the HTH and the HO-helix remains stable during all the time course of the dynamics (**Fig. 5b, 5c**). Apo and OM differ mainly in the fact that the Lever arm dynamics is quite reduced when OM is bound. Since OM allows fast rates of actin-driven P_i_ release, these simulations suggest that the actin-driven transitions that drive P_i_ release can occur without a large Lever arm swing.

In contrast, the simulations with Mava bound show that movement of the Switch elements around the P_i_ occurs on the first ns of the simulation, prior to a movement of Mava from its initial position. This movement closes the backdoor, which remains in this closed position throughout the simulation thanks to electrostatic bond compensation (**Fig. 5f, Supplementary Fig. 10, Supplementary Movie 4**). After rotation of the F465 side chain (at 11 ns), the communication between the Switch-2 and the Wedge is quickly lost around 36 ns (**Fig. 5f, Supplementary Fig. 10, 11, Supplementary Movie 4**). This loss of anchoring induces large divergence in the L50 orientation compared to what is explored when OM or no drug are bound. The L50 subdomain behaves as if it is no longer maintained while Switch-2 remains in a strongly closed position, trapping P_i_ (**Supplementary Movie 2, 3, 4**). Close analysis of the simulations suggests that specific interactions of Mava with ^HW^H666 and with the β1-β2 strands of the Transducer could create new constraints, central for altered communication between the drug-binding site and the active site. Thus, the simulations demonstrate that the presence of Mava alters the energy landscape of myosin conformations explored. The new allosteric communication within the head not only maintain P_i_ trapped but they also translate into divergent movements of the outer part of the internal myosin cleft. Thus more freedom for the L50 subdomain significantly alters the actin-binding interface, which deviates from the PPS-like conformations explored when OM is bound or no drug is present (apo). This is confirmed by the relative dynamics of the HTH compared to the HO-helix (**Fig. 5b, 5c**). While the HO-helix remains stable, the position of the HTH rapidly deviates from its position, indicating a strong alteration of the actin-binding interface. Thus, Mava-bound substrates are inappropriate for efficient starting point of the powerstroke (**Fig. 5f**).

Additionally, we observe rapid pocket exploration by the small Mava molecule, which can lead to its release in the solvent in one of the simulations, and allows Mava to occupy new internal pockets which are closed in other simulations. Mava indeed takes opportunities of movements of the β1-β2 strands and the linker preceding the HE helix to fit in new pockets (**Supplementary Fig. 10, Movie 3, Supplementary Movie 4, 5, 7**), thus stabilizing atypical conformations of the motor inadequate for Pi release or actin binding. This strongly differ from the much reduced dynamics observed for the longer OM molecule that fits in a closed drug-binding pocket stabilized by polar interactions made between residues of the N-term and Converter subdomains (**Supplementary Fig. 9, Movie2, Movie 3, 6, 7**).These differences in the mobility of Mava compared to OM is reflected in the Kd calculations over the time course of the simulation (**Supplementary Fig. 11**). While the calculated Kd of OM oscillates around 1 µM (average Kd 1.30 µM) and remains stable, that of Mava goes through larger oscillations due to the occupation of diverse pockets (average Kd 3.33 µM).

### A proposed mechanism for Mava

The structures combined to the dynamics allow to propose a mechanism of action explaining how the two drugs, occupying the same pocket can have distinct effects. The two drugs maintain the Lever arm up, but have **(i)** different dynamics within the pocket and **(ii)** distinct allosteric effects due to their differences in shape and structure.

The small and bulky Mava fits in a slightly wider pocket compared to the elongated OM. It maintains the Converter further away from the motor domain. Since it is only composed of two cycles, Mava does not establish interactions with the Relay, yet it explores pockets that are found near the first three strands of the Transducer (**Supplementary Fig. 10, Supplementary Movie 4, 6, 7**). Thus, it does not exert restraints on the Relay, explaining why the position of this connector deviates during the simulation, allowing both the Wedge and Switch-2 to loose communication and reach uncanonical positions. This has two consequences: **(i)** the backdoor maintains a closed conformation as Switch-2 movements greatly differ from what is observed for Apo myosin ; **(ii)** the actin binding interface is altered. Mava binding thus results in “incompetent PPS” states that interact poorly with actin and thus cannot release P_i_ readily. This is fully consistent with *in vitro* data demonstrating that when Mava is bound to β-cardiac myosin, the P_i_ release rate is slowed with and without actin and the affinity of the myosin head for actin is reduced^24,14^.

In contrast, OM is elongated and able to interact with the Relay helix. It consequently not only stabilizes the Lever arm up, but also exerts restraints on the Relay position. In contrast to Mava, it favors states with a primed Lever arm that are also able to interact with actin readily (**Supplementary Fig. 9, Movie 3, 5, 7**).

### Effects on the formation of the sequestered state

Since reports indicate that OM destabilizes the sequestered state while Mava seems to destabilize it^18,19,21^, we examined whether the high-resolution structures reported here were providing insights on this potential opposite effect. In cardiac IHM, both heads adopt a PPS state^7^, a structural state favored when ADP.Pi is bound, and which is largely similar to those favored by binding of OM or Mava. We next examined the precise Transducer conformations as well as its relative positioning to that of the Converter. The surface of these two structural elements of the motor domain indeed constitutes the free head (FH) mesa – a flat region recognized by the blocked head (BH)^10,7^. Importantly, the modulator bound in the OM/Mava pocket plays a key role for the relative positioning of the Transducer and the Converter. We demonstrate that when OM is bound, the surface differs greatly from that required for forming the FH Mesa due to the narrow size of the pocket and the change in the Transducer conformation (Fig. 6a). In contrast, the same surface of the Mava-bound structure does not differ significantly compared to that found in the apo structure and could thus allow interactions found between heads in the IHM. Quite interestingly, in the PPS-S1-Mava structure, the two molecules in the asymmetric unit interact with each other by forming interactions that are similar to those found between heads in the IHM (Fig. 6b). In summary, the Mava conformation does not differ from that required to form intramolecular IHM interactions mediated by the BH or the FH heads^7^ (Fig. 6c). In contrast, the OM-bound structure indicates that the activator changes the conformation of the Transducer, providing a first clue to explain why OM destabilizes the IHM.

**Figure 6.**
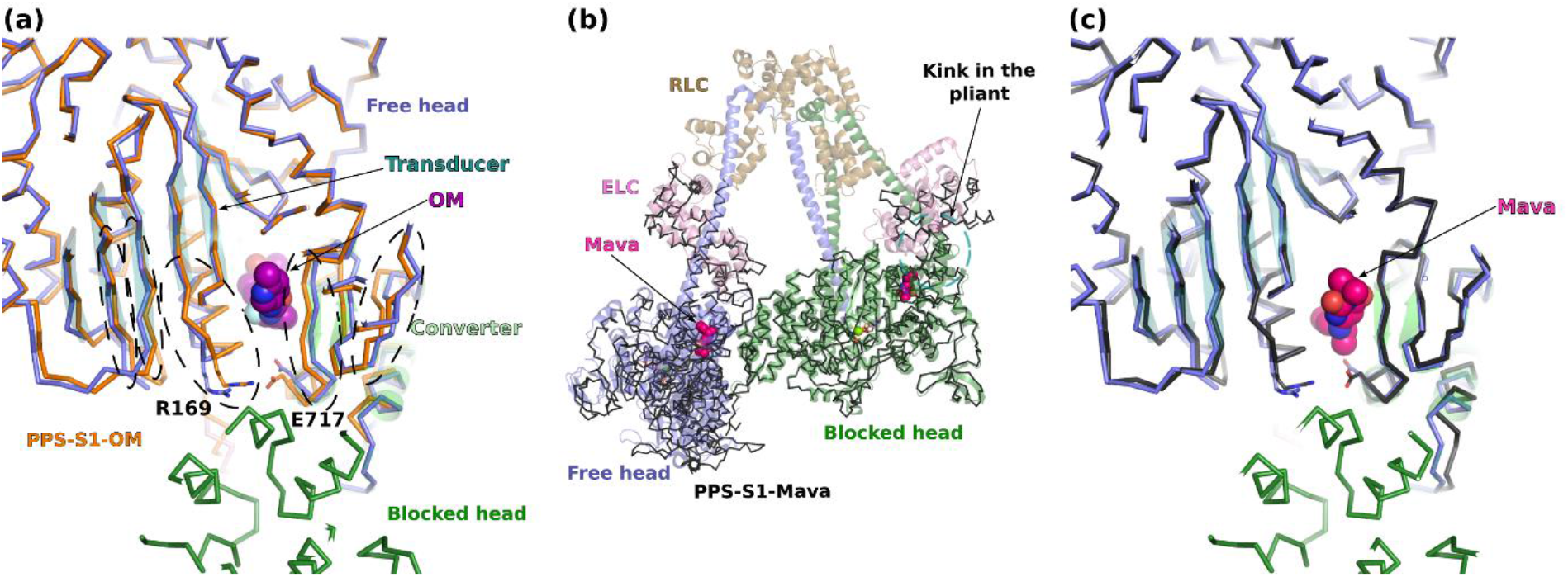
– Effects of Mava and OM on the β-cardiac myosin sequestered state. **(a)** Superimposition of the cryo-EM β-cardiac IHM structure (cartoon, colored, PDB code 8ACT ^7^ on the S1 structure of β-cardiac myosin complexed to OM (PDB 5N69, ^25^. Superimposition is done on the motor domain of the blocked head (BH) (residues 1-707). The presence of OM induces several differences in the Transducer and in the Converter (indicated in dashed lines). **(b)** Superimposition of the β-cardiac IHM structure on the two molecules of the asymmetric unit of PPS-S1-Mava (ribbon, in black). The superimposition was performed on the head-head interface (residues 329-447 of the BH and residues 498-518, 708-780 on the FH, rmsd 0.5 Å on 112 Cα). The structures are highly superimposable. **(c)** Superimposition of a molecule of the asymmetric unit of PPS-S1-Mava on the BH (residue 1-707) of the β-cardiac myosin IHM

## Discussion

Small-molecules able to modulate the force produced by myosins are today the best hope for treatments against acute heart failure and inherited heart diseases such as cardiomyopathies^42^. Amongst these treatments, the activator *Omecamtiv mecarbil* (OM) and the inhibitor *Mavacamten* (Mava) are the most advanced: OM is in phase 3 clinical trials against heart failure^13^ and Mava was approved by the FDA under the name of CAMZYOS^TM^ to treat hypertrophic cardiomyopathies^43^. While such small-molecules will be used as treatments, their precise mechanism of action must be revealed. This requires understanding of how they allosterically influence motor function. In addition, this information is of invaluable interest for the conception of specific myosin modulators that can treat a large range of myosin diseases^11^.

In this work, we combined high-resolution X-ray structures and molecular dynamics to elucidate the molecular mechanisms of OM and Mava, two molecules with antagonistic effects on force output of the heart^14,24,17^. We reveal that Mava occupies the same pocket as OM, a binding site that only forms in states of the motor in which the Lever arm is primed. Local changes near the drug-binding site differ when the Apo, OM and Mava structures are compared and these small differences can be transmitted to the Transducer and the Lever arm which are key-elements of the allosteric communication in myosin motors^6^ .

Detailed analysis coupled to molecular dynamics reveal how OM favors the exploration of structural states that are similar to the apo motor, while Mava triggers exploration of “incompetent” PPS states in which the actin interface and coupling within the motor are perturbed, consistent with the previously observed decrease of P_i_ release rate^24^. Single-molecule experiments demonstrated that OM is compatible with actin binding, although it maintains the Lever arm primed^26^, acting as a suppressor of the powerstroke. Our results are fully consistent with this data, as the cohesive action of OM around its binding site can help maintain the Lever arm primed when it occupies the pocket. The powerstroke would require release of OM from this binding site. In contrary, Mava slows actin binding since the actin-binding interface is altered due to uncoupling of the myosin subdomains. This particularly affects the internal cleft, whose conformation and dynamics greatly impact the actin binding interface. Interestingly, we observed that closure of Switch-2 could also participate in reducing the P_i_ release rate^24,14^ by closing the backdoor. In light of the structural states explored, while OM increases the number of competent heads able to interact with actin filament and produce force, Mava reduces this number by trapping the heads in an incompetent ensemble of pre-stroke states that cannot efficiently interact with actin and initiate force production (Fig. 7). Thus, all atom dynamics simulations provide essential insights on how binding of small-modulators alter the allosteric communication within a myosin motor, in particular by altering the conformational ensembles explored by the motor domain and decoupling these movements from the priming of the Lever arm.

**Figure 7.**
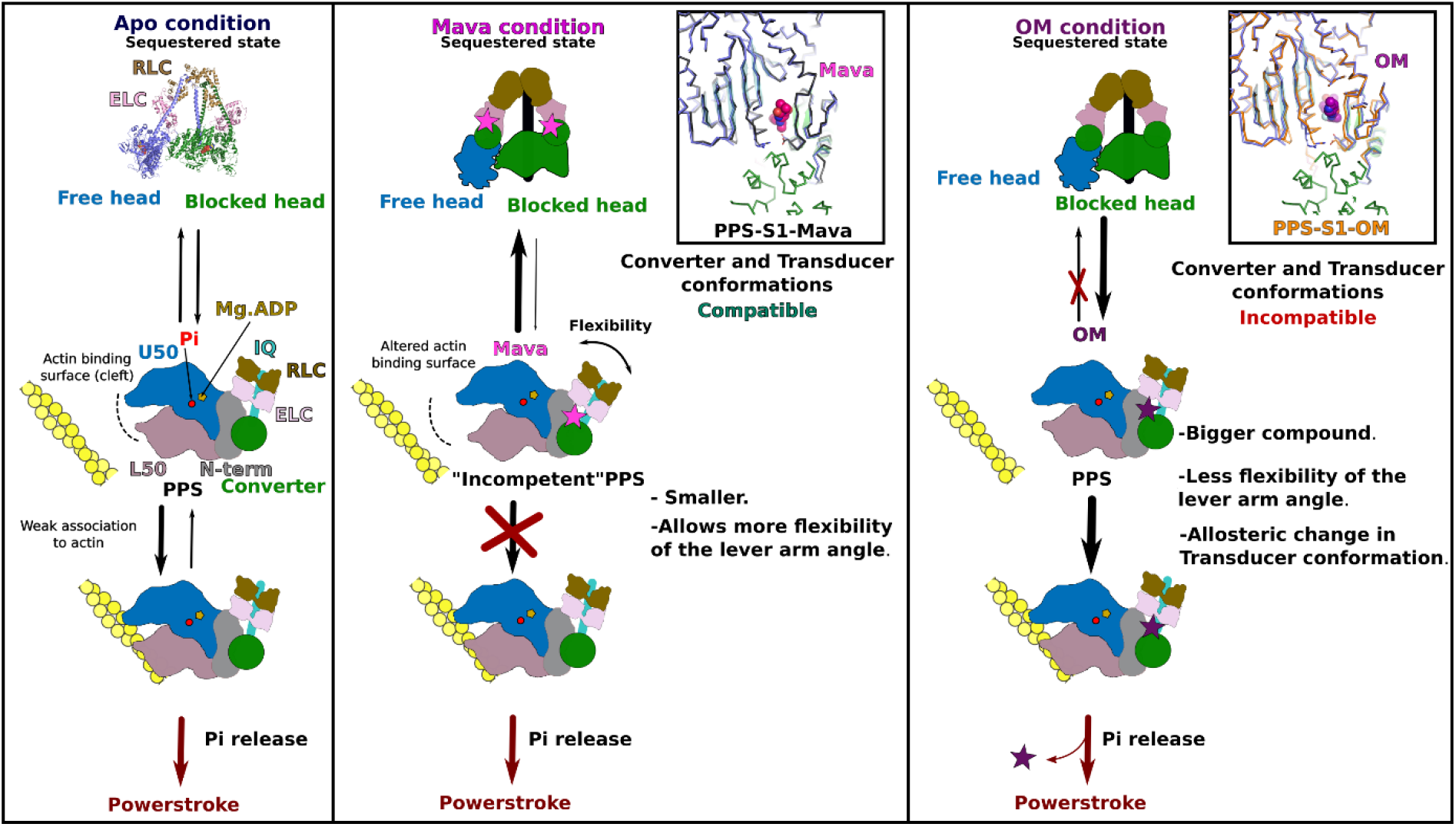
– OM and Mava mechanism of action. Schematic representation of the effects of both OM and Mava on **(i)** folding back to form the sequestered state and **(ii)** P_i_ release. **(Left)** The apo condition (without drug) allows the dynamic formation of the sequestered state (Interacting-heads motif, IHM) in equilibrium with heads available for association with actin. **(Center)** Mava alters the actin binding interface, generating an “incompetent” PPS that slows association with the actin filament. Mava-bound heads undergo flexibility in lever arm priming, a requirement for the formation of the head/head interfaces characteristic of the cardiac sequestered state^7^. In the zoom (inset), the cardiac IHM blocked and free head structures are compared with the Mava-bound crystal structure (thin black lines). **(Right)** When OM binds (thin orange lines), the conformation of the Transducer and the orientation of the Lever arm differ from what is found in cardiac IHM. PPS-OM is thus incompatible with the sequestered state. OM bound heads can efficiently bind F-actin and promote P_i_ release, yet the swing of the lever arm is affected as it requires destabilization of the OM binding site^26^.

Mava and OM have also been reported to have antagonistic effects on the sequestered auto-inhibited state of cardiac myosin^19,20,21^. Thus, the two drugs also affect the ability of two heads to form the IHM interface, although they bind in the same pocket. The high-resolution structure of β-cardiac myosin IHM without drug bound^7^ provides the precise description of the head/head interfaces. The FH Mesa corresponds to the surface of the FH head involved in interactions with the BH head, it involves the Transducer and the Converter. The OM structure indicates why the surface required to bind to the BH head is not compatible with the formation of the IHM. Closure of the pocket around OM and allosteric changes on the Transducer both impose important changes in the FH Mesa that can prevent the formation of the IHM (Fig. 7). In the Mava-bound structures, no allosteric alteration of the Transducer conformation is observed and all atom simulations indicate that mobility exists in the drug binding pocket that might be used to form the interactions required for IHM stabilization. Consistently, FRET measurements have indicated that Mava increases the number of heads forming the IHM by ∼4%^21^. Interestingly, this amount is one order of magnitude less than previously reported when a crosslinked version of Mava has been used^19^. Currently, a high-resolution structure of the IHM with Mava bound is lacking. The medium resolution structures (∼6 Å at most^9,8^) cannot directly assess whether the modulators are bound. Given the effects of Mava on the motor domain conformation observed in dynamics, we cannot exclude that the drug induces differences in the IHM conformation that are not visualized in the filament structures due to the limit in resolution. Future studies of a high-resolution Mava-bound cardiac IHM is thus required. Importantly, the effects of Mava on the stabilization of the IHM seems modest^21^. Its 10-fold higher effect on mant-nucleotide exchange – often called SRX measurements^21^ – indicates that the drug efficiently inhibits product release. Uncoupling within the motor head as seen in our all-atom simulations provides insights on why the presence of Mava induces slower binding to F-actin, as well as slower release of bound nucleotide.

To conclude, the results presented here provide the blueprint of the inhibition vs. activation of β-cardiac myosin by two different molecules targeting the same pocket. It illustrates how distinct restraints applied on the N-term, the Converter and some connectors such as the Relay can govern distinct allosteric behavior in the myosin motor. Modulators with distinct scaffolds can bind and stabilize this pocket differently indicating how the plasticity would regulate drug affinity as well as allosteric effects in motor function. Interestingly, this pocket also affects myosin regulation by controlling the IHM formation. All atoms molecular dynamics grounded on high-resolution structures is of prime interest for understanding of how force is generated and how myosin modulators can be conceived to regulate force generation in the context of human diseases. Cardiomyopathies can result from distinct mutations and the description of the OM/Mava pocket opens the door to exploring how to modify the potency of these derivatives for customizable effects on force production depending on the pathological phenotype to be corrected. Such a rational approach will also be of interest for other myosins since modulators are being studied to target other myosin-associated pathologies such as spasticity by targeting SkMyo2^44^; malaria by targeting myosin A^35,36,45^, or asthma by targeting SmMyo2^46^.

## Data availability

The atomic models and the structure factors of PPS-MD-Apo, PPS-S1-Mava, PPS-MD-Mava and PPS-S1-OM are available at the PDB under the accession numbers 8QYP, 8QYQ, 8QYR and 8QYU respectively.

## Acknowledgements

The authors greatly acknowledge Dr. James Hartman (Cytokinetics inc.) for providing purified β-cardiac myosin S1; Dr. Catherine Guillou and Dr. Thibault Bayles (Institut de Chimie des Substances Naturelles, ICSN, UPR2301, Gif-sur-Yvette, France) for providing the Mavacamten derivatives that help identifying the initial crystallization conditions ; the beamline scientists of PX1 and PX2A (SOLEIL synchrotron) for excellent support during data collection. The A.H. team is part of the Labex Cell(n)Scale ANR-11-LBX-0038 and IDEX PSL, which is part of the Initiatives of Excellence of Université Paris Sciences et Lettres (ANR-10-IDEX-0001-02-PSL). This work was supported by the CNRS, grants to A.H. from AFM 21805, FRM DCM20181039553, NIH RM1GM131981-01 and ANR-21-CE11-0022-01.

## Authors and contributions

AH designed and directed the research. JRP and AH conceived and planned the experiments and were involved in project administration. GT performed the synthesis of Mavacamten. Protein crystallization was performed by JRP with the help of AD. Data collection and processing was performed by JRP with the help of CK. JRP and CK performed model building and refinement. Molecular dynamics experiments were run by DA with the help of SR. SR provided access to the calculators and helped to set up the calculations. AWS and HJK designed and supervised the chemical synthesis of Mavacamten and analyzed the synthesized compounds. Formal analysis and validation of the results was performed by DA, JRP and AH. JRP, DA and AH wrote the manuscript with the help SR and all authors reviewed it. AH provided funding.

## Competing interests

AH receives research funding from Cytokinetics and consults for Kainomyx. All other authors have no competing interests.

## Supplementary Figures

**Supplementary Figure 1.**
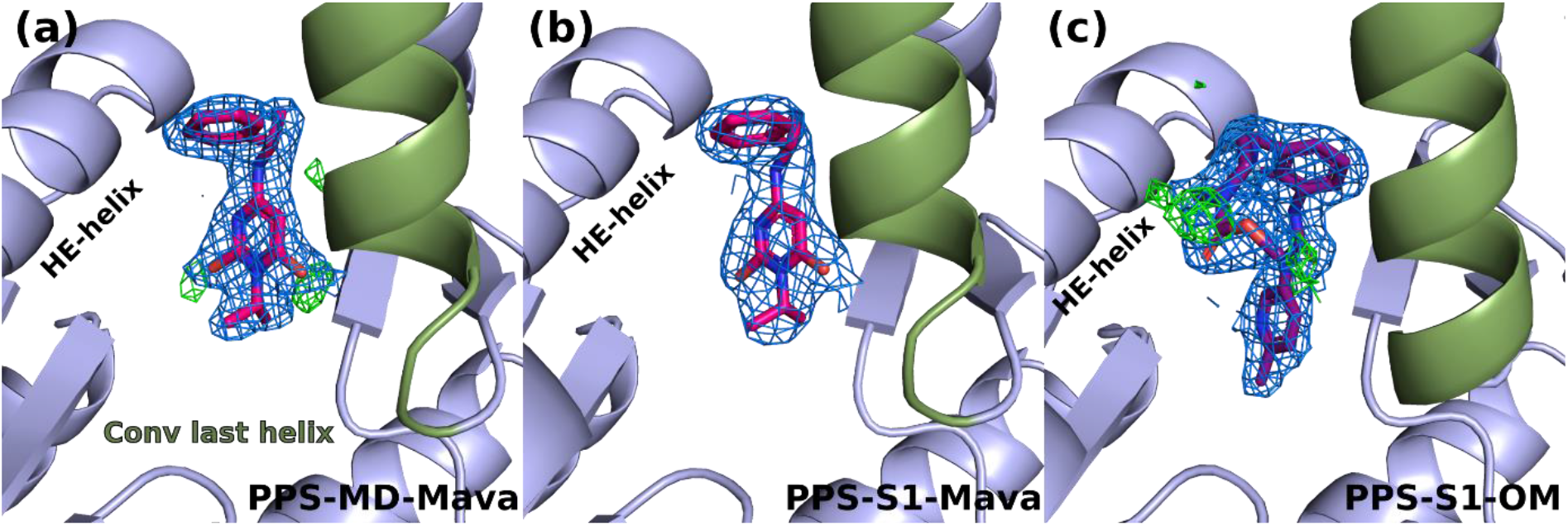
– The drugs are defined without ambiguity in the electron density maps. The three panels present the electron density maps corresponding to the drugs bound in the three β-cardiac myosin structures: **(a)** PPS-MD-Mava; **(b)** PPS-S1-Mava and **(c)** PPS-S1-OM. The 2Fo-Fc map contoured at 1 σ is represented in blue; the positive peaks of difference map Fo-Fc, contoured at 3 σ, are represented in green. The last helix of the Converter is colored in smudge green and labelled as “Conv last helix”.

**Supplementary Figure 2.**
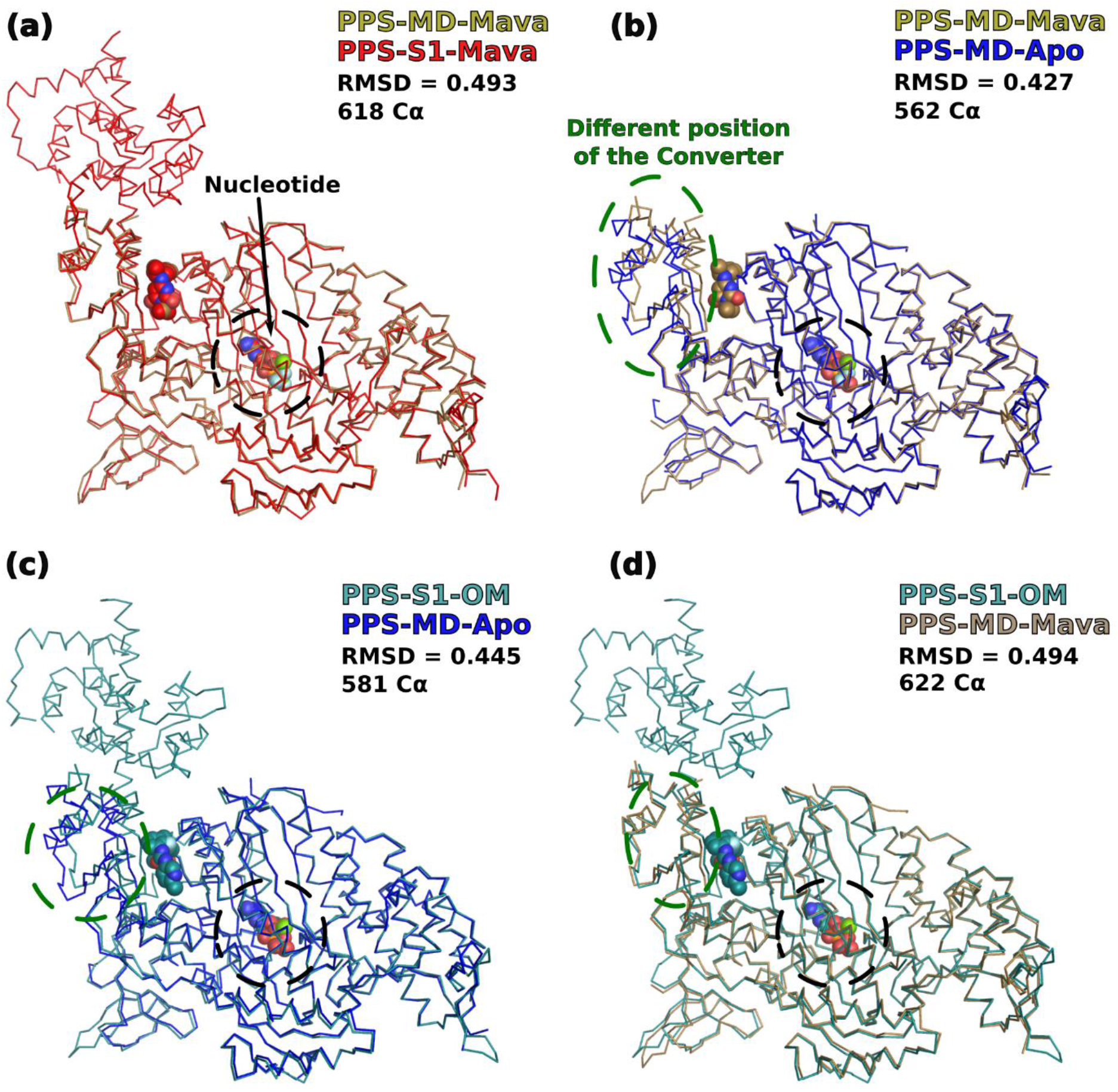
– Comparison of the different β-cardiac myosin PPS structures from this work. **(a)** Superimposition of the PPS-MD-Mava and PPS-S1-Mava chain A; **(b)** PPS-MD-Mava and PPS-MD-Apo; **(c)** PPS-S1-OM chain A and PPS-MD-Apo; **(d)** PPS-S1-OM chain A and PPS-MD-Mava. The RMSD on Cα is indicated as well as the colors used for each of these structures. The alignment was performed using a selection of the Cα in the motor domain (residues (1-780)). The drugs and the nucleotide are represented in spheres. The drug position is contoured in black with dashed lines. When the position of the Converter differs significantly between the structures compared, the Converter position is contoured with green dashed lines.

**Supplementary Figure 3.**
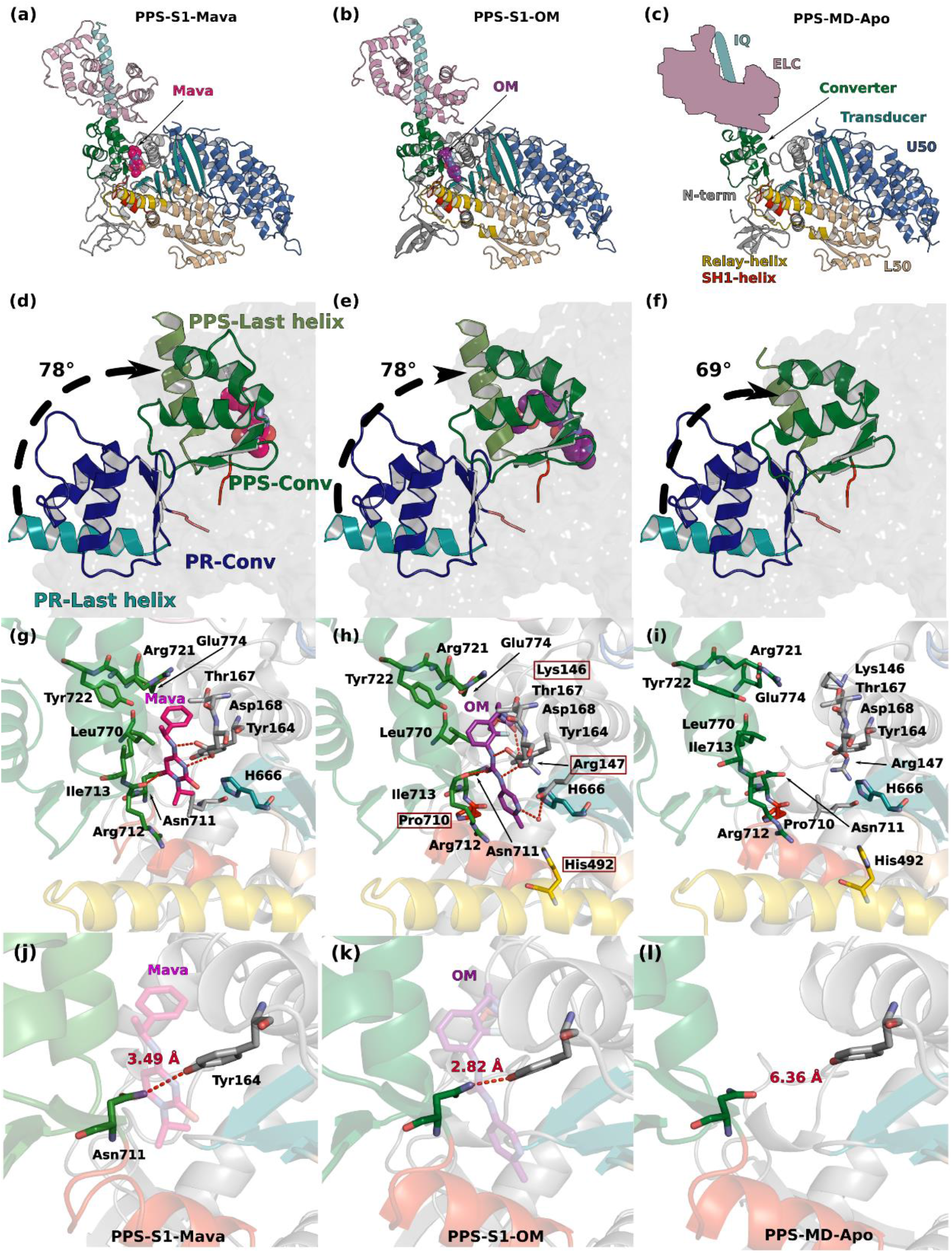
– OM and Mava have distinct effects on the priming of the Lever arm. **(a), (b), (c)** Overall structures of PPS-S1-Mava, PPS-S1-OM and PPS-MD-Apo. **(d)**, **(e)** and **(f)**, Comparison of the priming of the Lever arm, while the structures are superimposed on the first 707 residues of the motor domain. The angle is measured between the PPS and the post-rigor structure (PDB code 6FSA^1^) using the position of the last helix of the Converter (769-780). **(g)**, **(h)**, **(i)** Zooms on the OM/Mava binding pocket. **(j)**, **(k)**, **(l)** Effects of drug binding on pocket size.

**Supplementary Figure 4.**
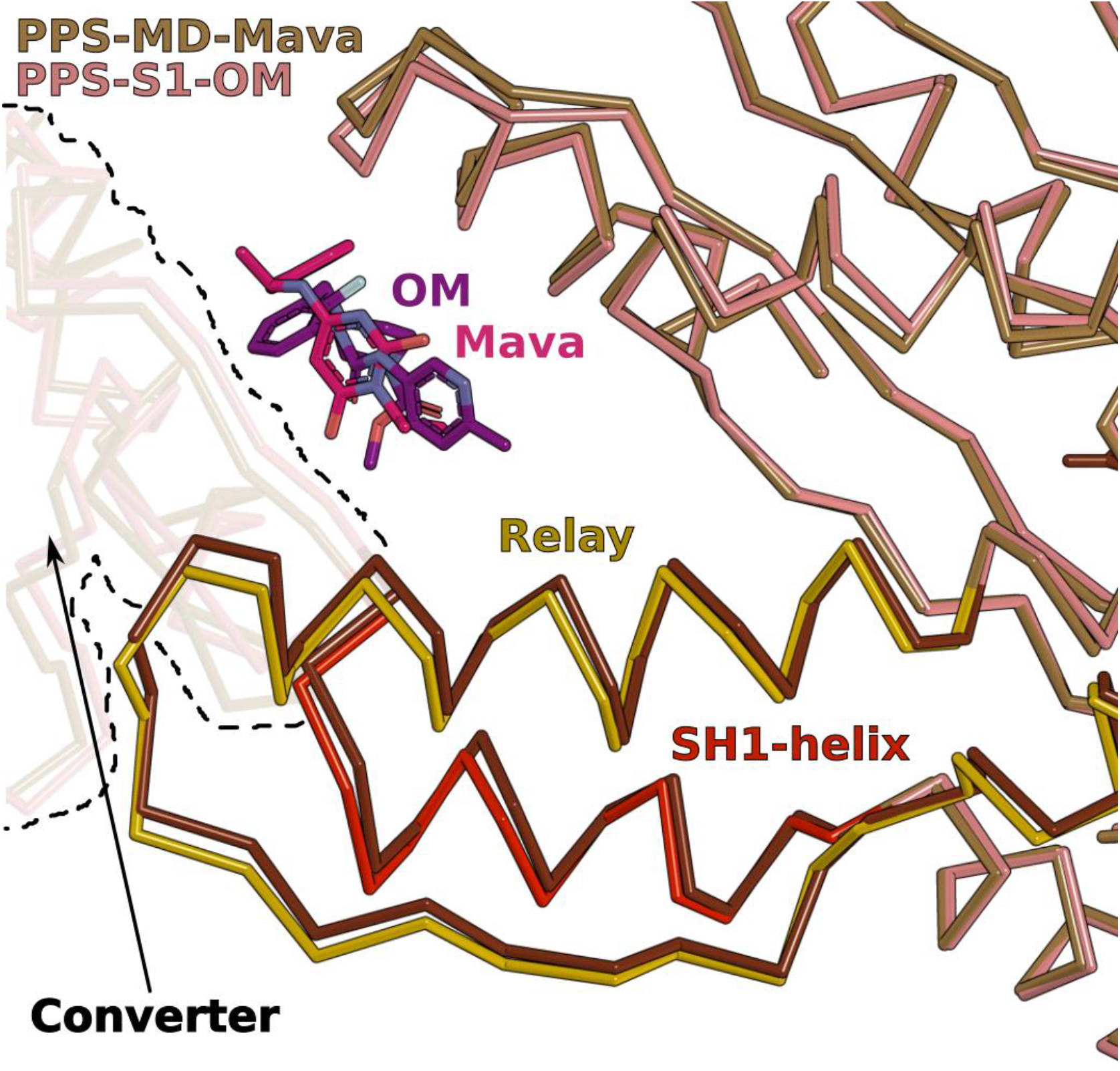
– Comparison of the pocket size when OM and Mava are bound. The PPS-S1-Mava and PPS-S1-OM structures are superimposed on the N-term subdomains. The position of the Relay and SH1-helices differ in the two PPS structures when OM or Mava are bound. Note also in the back how these differences correspond to different positions of the Converter in the two structures (contoured with black dashed lines).

**Supplementary Figure 5.**
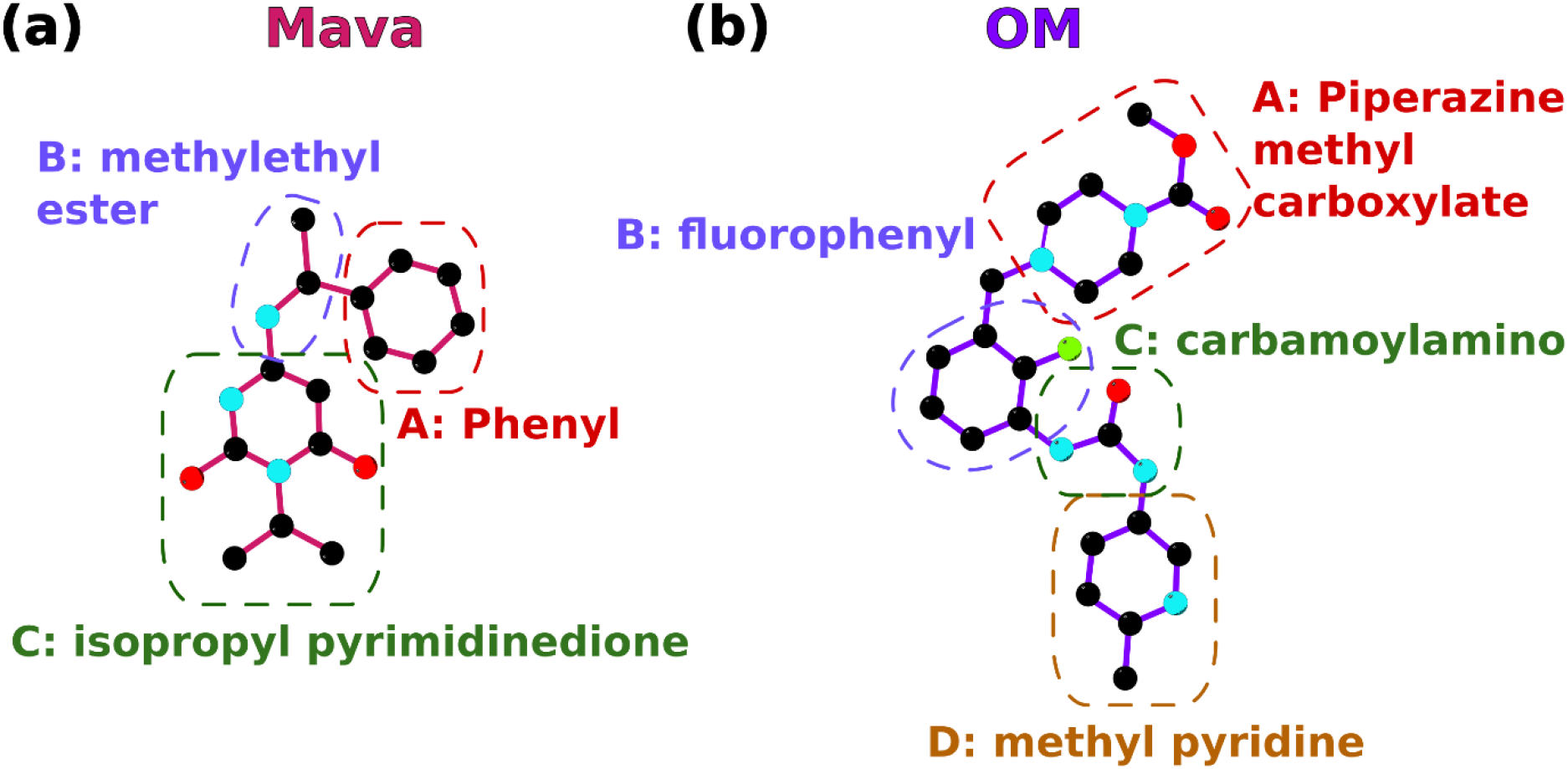
– The different chemical groups of Mava and OM. **(a)** Schematic view of the chemical groups of Mavacamten: B: phenyl, B: methylethyl ester; C: isopropyl pyrimidinedione. **(b)** Schematic view of the chemical groups of Omecamtiv mecarbil: A: piperazine methyl carboxylate; B: fluorophenyl; C: carbamoylamino; D: methyl pyridine.

**Supplementary Figure 6.**
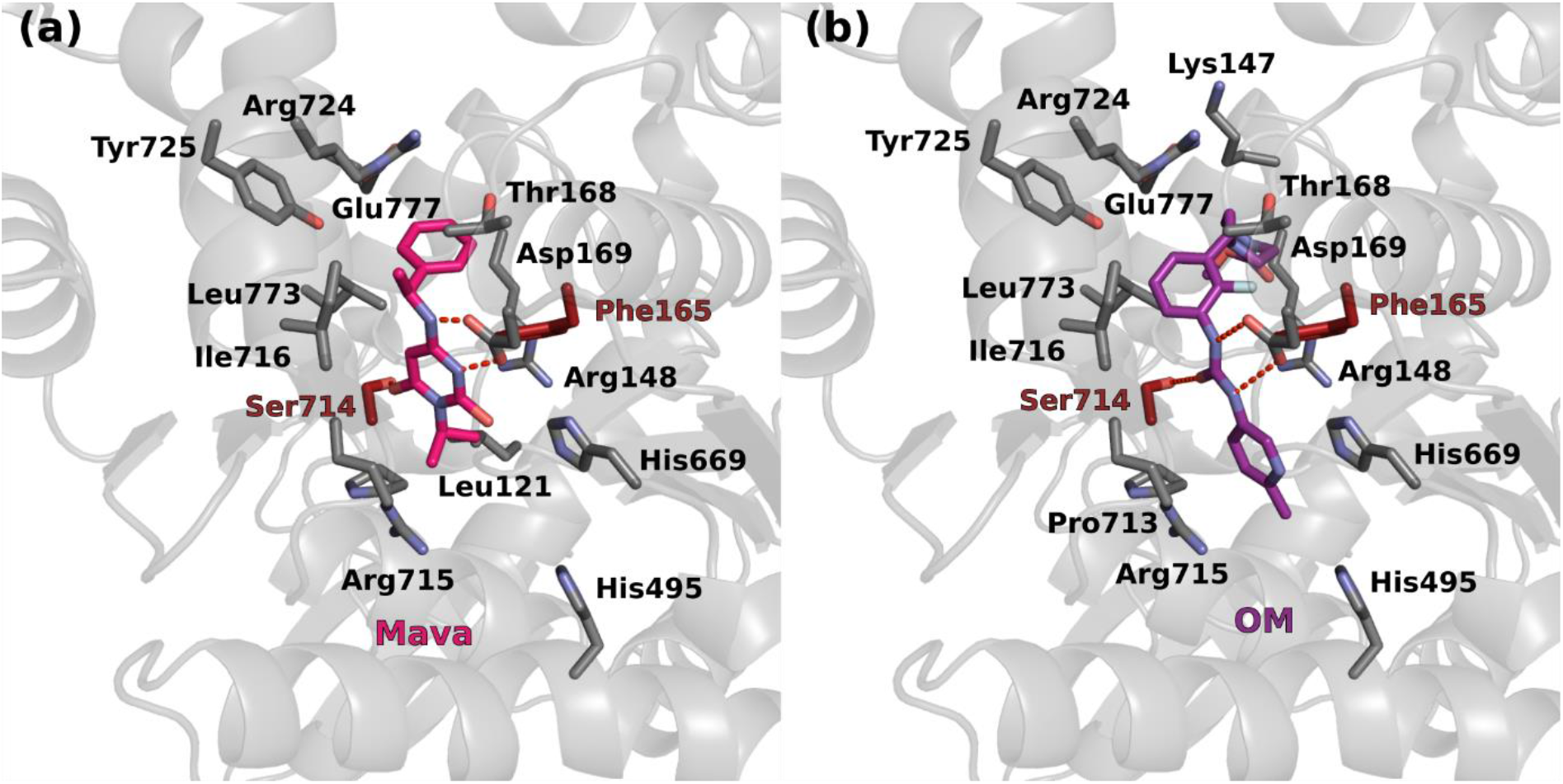
– Homology model of Oryctolagus cuniculus skeletal myosin 2 (SkMyo2_Oc) in PPS bound to (a) Mava or (b) OM. The PPS-S1-OM structure was used as a template for the modelling. Different residues are colored in red.

**Supplementary Figure 7.**
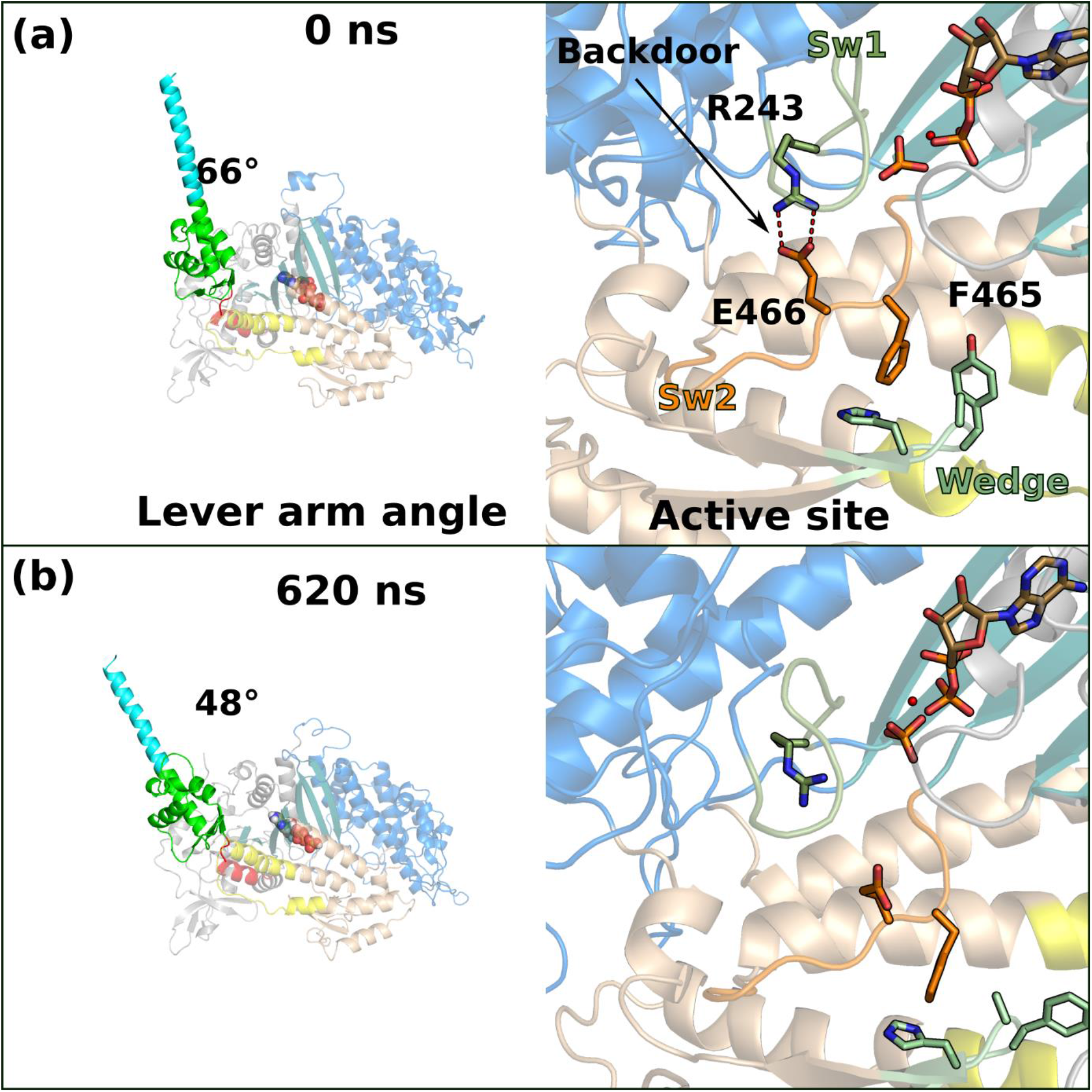
– All atom molecular dynamics of the S1 fragment in the apo form. The consequences on the Lever arm angle (left) and on the active site (right) are represented.

**Supplementary figure 8.**
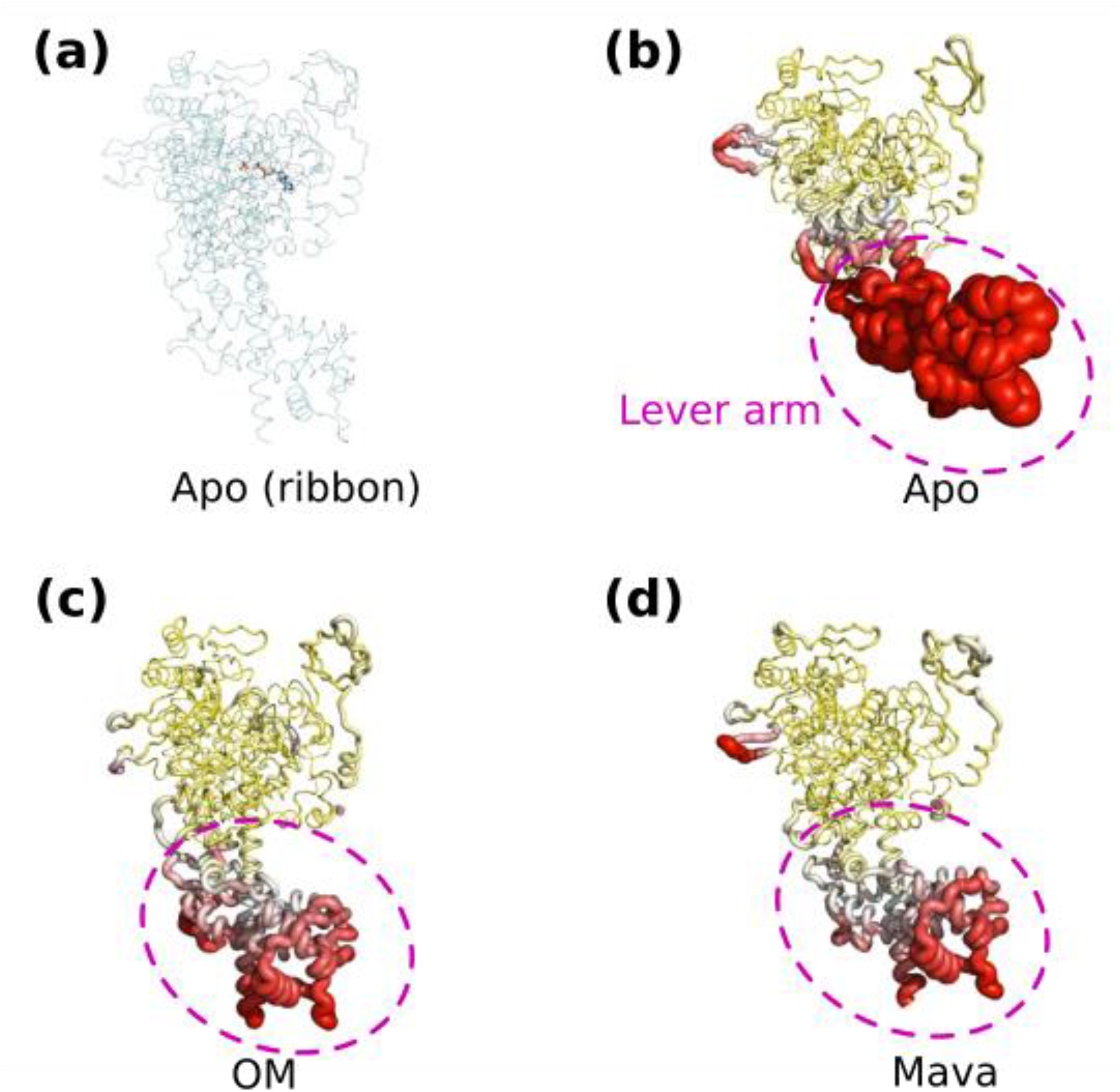
– Patterns of mobility for the different conditions starting from a similar conformation (PPS) **(a)** Apo S1-PPS + ADP.Pi hybrid model as visual reference. **(b)** “Putty representation” of the 730 ns trajectory of the Apo form. **(c)** “Putty representation” of the 1050ns trajectory of the OM form. **(d)** “Putty representation” of the 1050ns trajectory of the Mava form. All four subsets are depicted in the same referential. All “Putty representations’ (using the ‘psico module’ PyMOL) are colored in yellow–white–red gradient with rmsd ranging from 0 to 9 Å. (Note that the maximum deviation for the Apo, OM and Mava MD simulations is 23.2 Å, 13.3 Å and 14.3 Å, respectively).

**Supplementary Figure 9.**
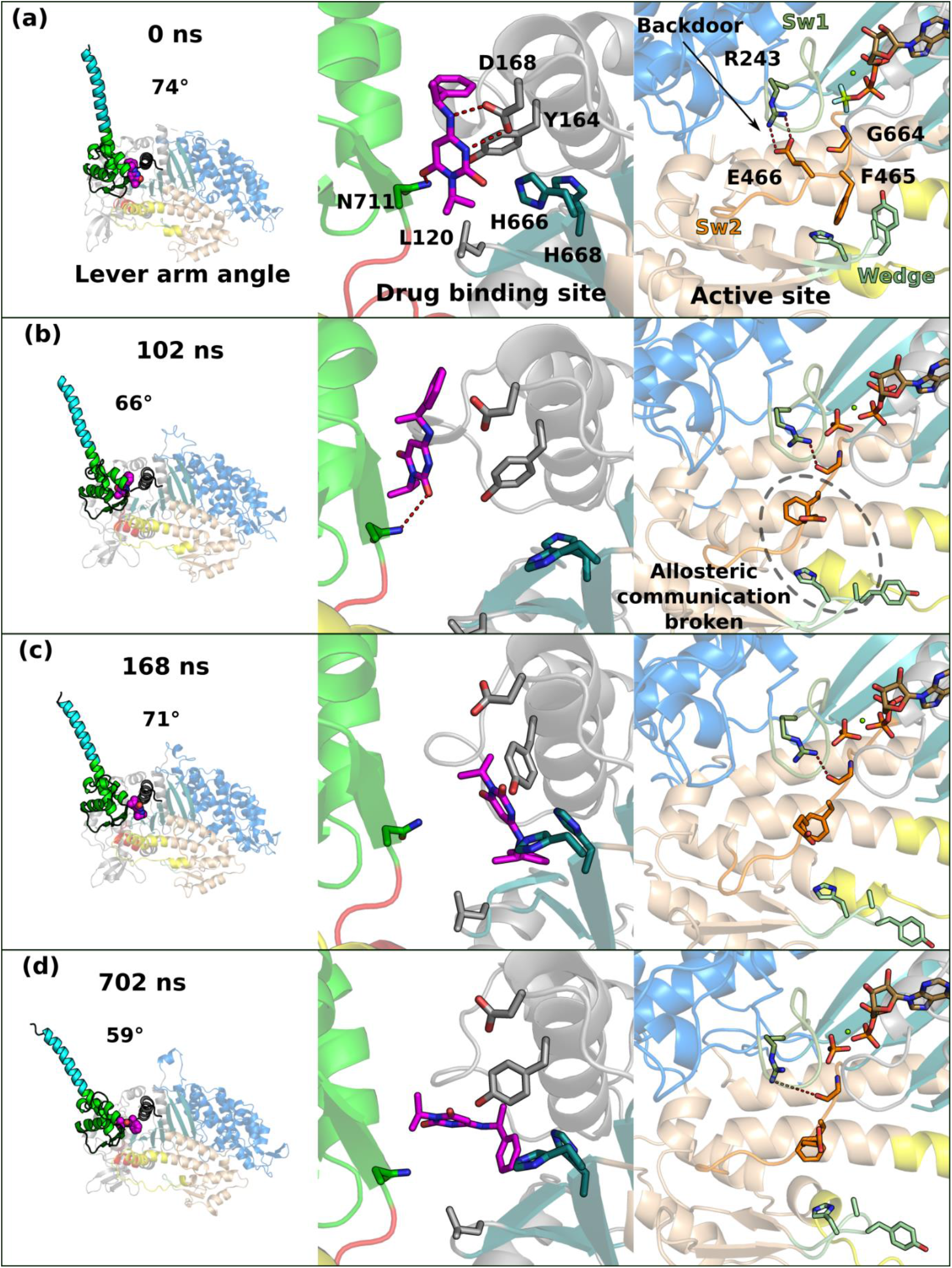
– All atom molecular dynamics of the S1 fragment bound to Mava. **(a), (b), (c)** and **(d)** Representative times at which Mava occupies distinct positions in the pocket. The side chain of key residues are represented as sticks. The consequences on the Lever arm angle (left), on the pocket (center) and on the active site (right) are represented.

**Supplementary Figure 10.**
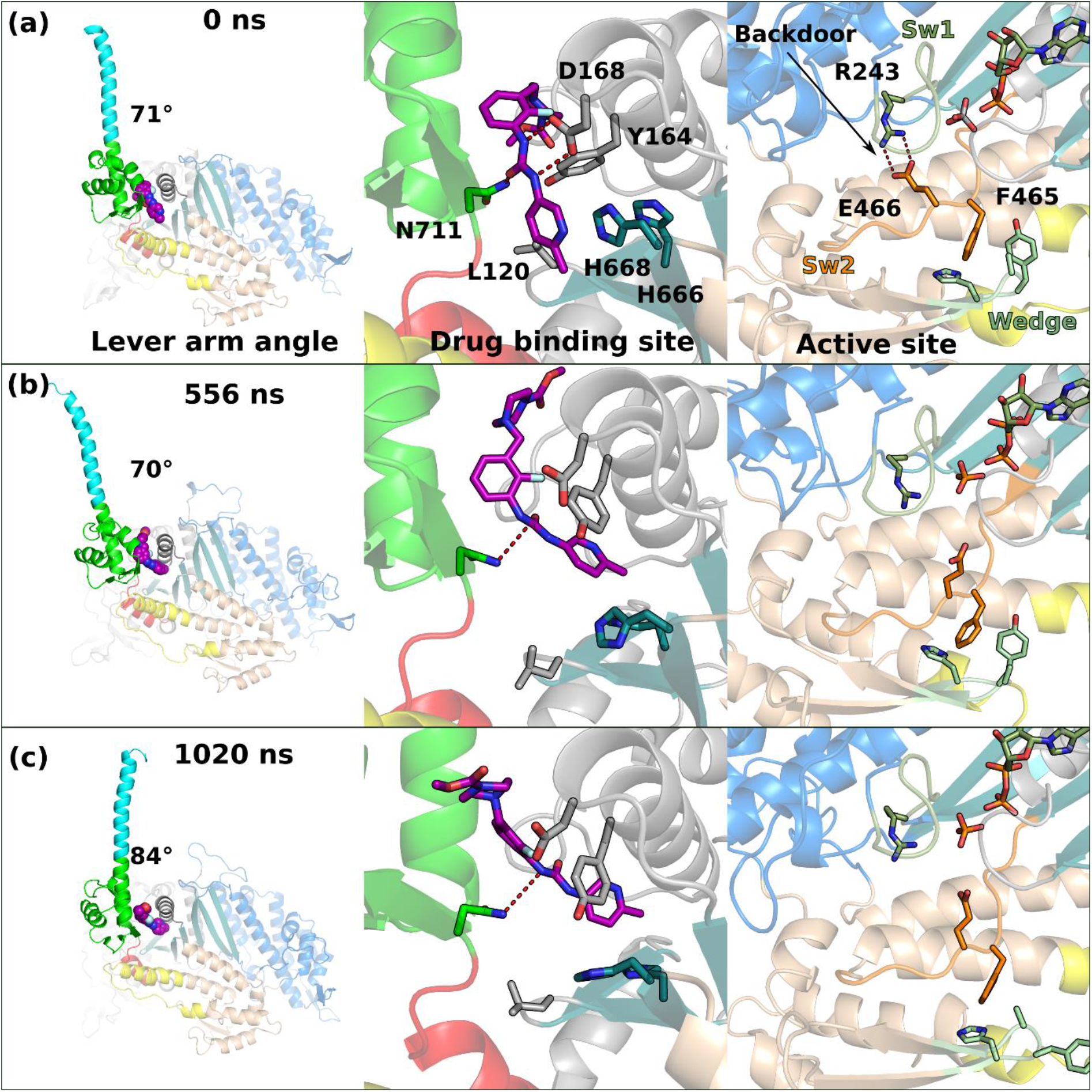
– All atom molecular dynamics of the S1 fragment bound to OM. **(a)**, **(b)** and **(c)** Representative times at which OM occupies distinct positions in the pocket. The side chain of key residues are represented as sticks. The consequences on the Lever arm angle (left), on the pocket (center) and on the active site (right) are represented.

**Supplementary Figure 11.**
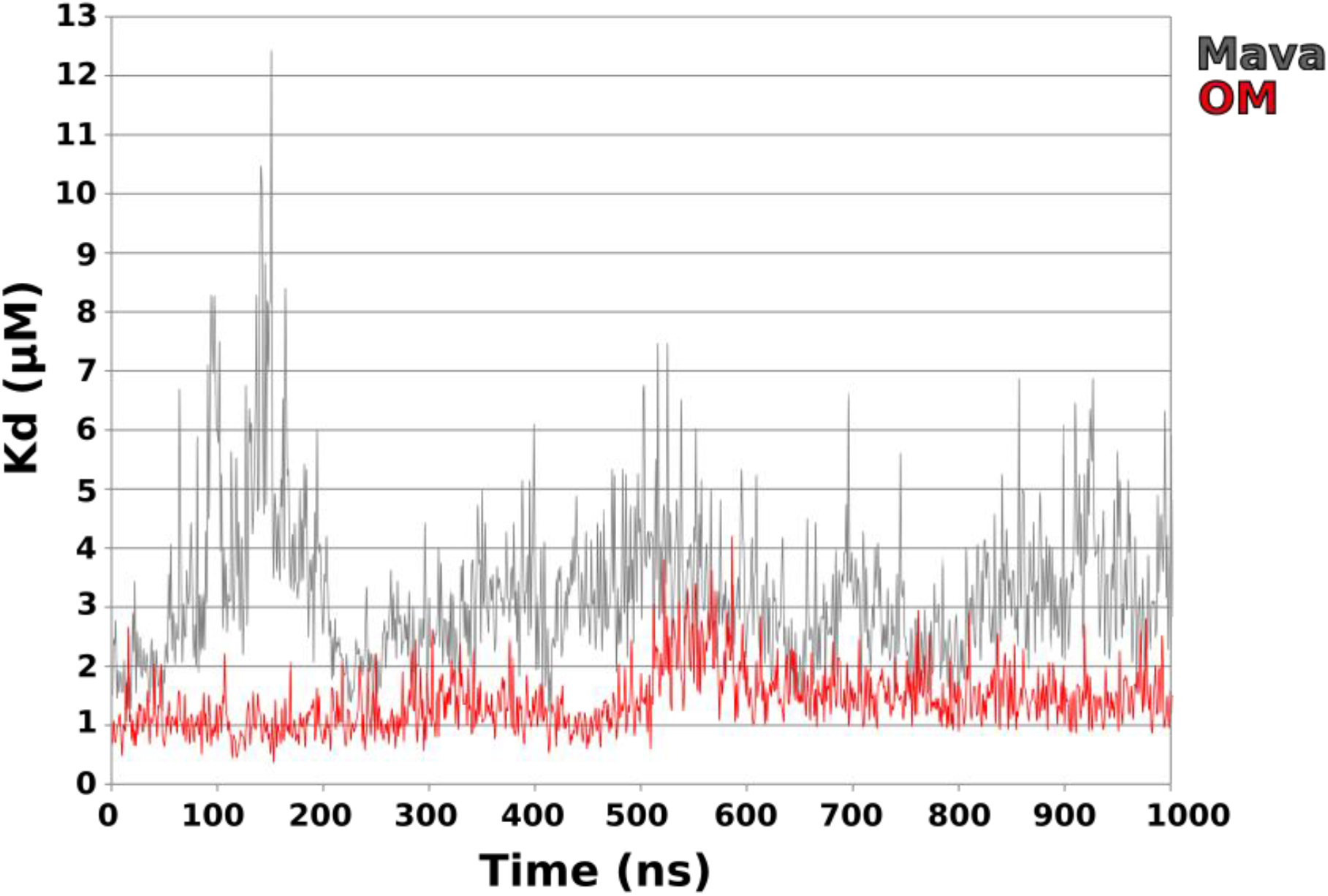
– Evolution of the dissociation constant (Kd) of OM and Mava during the time course of the simulations. The binding constant was derived from the free energy calculations computed with PRODIGY-LIG^2,3^. During the OM MD simulation, the calculated Kd oscillates around ∼1 µM. During the Mava MD simulations, the calculated Kd of the more mobile Mava goes through larger variations. Despite this variability, the average calculated Kd of Mava is 3.3 µM.

**Supplementary Table 1.** – Data collection and refinement statistics (molecular replacement)

|  | PPS-MD-Apo | PPS-S1-OM |
| --- | --- | --- |
| **Data collection** |  |  |
| Space group | P4<sub>1</sub>22 | P2<sub>1</sub>2<sub>1</sub>2<sub>1</sub> |
| Cell dimensions |  |  |
| *a*, *b*, *c* (Å) | 94.397, 94.397, 219.293 | 97.996, 122.503, 187.463 |
| $\alpha$, $\beta$, $\gamma$ (°) | 90.000, 90.000, 90.000 | 90.000, 90.000, 90.000 |
| Resolution (Å) | 94.397-2.759 (3.033-2.759) | 102.547-1.928 (2.128-1.928) |
| *R*<sub>merge</sub> (all I+ & I-) | 0.273 (2.413) | 0.153 (1.267) |
| *R*<sub>merge</sub> (within I+ & I-) | 0.296 (2.355) | 0.139 (1.164) |
| Number of observations (total) | 522201 (26600) | 473345 (30533) |
| Number of observations (unique) | 20166 (1009) | 97819 (4891) |
| *I* / σ*I* | 10.8 (1.5) | 7.9 (1.5) |
| Completeness (Spherical) (%) | 76.3 (15.8) | 57.6 (11.3) |
| Completeness (Ellipsoidal) (%) | 94.4 (64.2) | 93.9 (73.7) |
| Redundancy | 25.9 (26.4) | 4.8 (6.2) |
| CC<sub>1/2</sub> | 0.999 (0.710) | 0.991 (0.519) |
| **Refinement** |  |  |
| Resolution (Å) | 86.73-2.759 (2.858-2.759) | 22.58-1.962 (2.032-1.962) |
| No. reflections | 20165 (190) | 92500 (636) |
| *R*<sub>work</sub> / *R*<sub>free</sub> | 0.2010/0.2595 | 0.1874/0.2355 |
| No. atoms | 5723 | 15636 |
| Protein | 5687 | 14699 |
| Ligand/ion | 33 | 168 |
| Water | 3 | 769 |
| *B*-factors | 77.73 | 38.62 |
| Protein | 77.84 | 38.69 |
| Ligand/ion | 62.35 | 32.02 |
| Water | 45.18 | 38.60 |
| R.m.s. deviations |  |  |
| Bond lengths (Å) | 0.016 | 0.048 |
| Bond angles (°) | 1.67 | 1.78 |

\*Number of xtals for each structure should be noted in footnote. \*Values in parentheses are for highest-resolution shell.
[AU: Equations defining various *R*-values are standard and hence are no longer defined in the footnotes.]
[AU: Ramachandran statistics should be in Methods section at the end of Refinement subsection.]
[AU: Wavelength of data collection, temperature and beamline should all be in Methods section.]

**Supplementary Table 2.** – Data collection, phasing and refinement statistics for PPS-S1-Mava and PPS-MD-Mava.

|  | PPS-S1-Mava | PPS-MD-Mava |
| --- | --- | --- |
| **Data collection** |  |  |
| Space group | P2<sub>1</sub> | P2<sub>1</sub>2<sub>1</sub>2<sub>1</sub> |
| Cell dimensions |  |  |
| *a*, *b*, *c* (Å) | 102.801, 147.977, 116.977 | 68.577, 95.684, 127.139 |
| $\alpha$, $\beta$, $\gamma$ (°) | 90.000, 91.625, 90.000 | 90.000, 90.000, 90.000 |
| Resolution (Å) | 91.635-2.607 (2.859-2.607) | 76.451-1.805 (1.986-1.805) |
| *R*<sub>merge</sub> (all I+ & I-) | 0.200 (1.212) | 0.185 (2.562) |
| *R*<sub>merge</sub> (within I+ & I-) | 0.177 (1.111) | 0.190 (2.498) |
| Number of observations (total) | 313583 (18434) | 1082068 (40777) |
| Number of observations (unique) | 69889 (3494) | 41812 (2083) |
| *I* / σ*I* | 7.4 (1.5) | 11.8 (1.3) |
| Completeness (Spherical) (%) | 65.8 (13.7) | 53.9 (10.9) |
| Completeness (Ellipsoidal) (%) | 93.6 (61.2) | 92.2 (64.6) |
| Redundancy | 4.5 (5.3) | 25.9 (19.6) |
| CC<sub>1/2</sub> | 0.985 (0.476) | 0.999 (0.467) |
| **Refinement** |  |  |
| Resolution (Å) | 84.40-2.610 (2.703-2.610) | 34.290-1.805 (1.869-1.805) |
| No. reflections | 69880 (530) | 41795 (75) |
| *R*<sub>work</sub> / *R*<sub>free</sub> | 0.185/0.226 | 0.183/0.225 |
| No. atoms | 14993 | 6258 |
| Protein | 14383 | 5763 |
| Ligand/ion | 125 | 71 |
| Water | 485 | 711 |
| *B*-factors | 63.44 | 38.50 |
| Protein | 64.29 | 38.44 |
| Ligand/ion | 43.10 | 40.44 |
| Water | 43.28 | 39.01 |
| R.m.s. deviations |  |  |
| Bond lengths (Å) | 0.014 | 0.015 |
| Bond angles (°) | 1.82 | 1.73 |

\*Number of xtals for each structure should be noted in footnote. \*Values in parentheses are for highest-resolution shell.
[AU: Equations defining various *R*-values are standard and hence are no longer defined in the footnotes.]
[AU: Ramachandran statistics should be in Methods section at the end of Refinement subsection.]
[AU: Wavelength of data collection, temperature and beamline should all be in Methods section.]

**Supplementary Table 3.** – Comparison of the interactions established by Mavacamten (Mava) and Omecamtiv mecarbil (OM)

| **Mava** | **β-cardiac myosin** | **OM** |
| --- | --- | --- |
|  | N-term**Arg147** | (A) |
|  | N-term**Lys146** | (A) electrostatic, carboxylate and εN-H |
| (A), (B), (C) stacking | N-term**Tyr164** | (A), (B), (C) electrostatic, Carboxylate, water, hydroxy |
| (B), (C) electrostatic | N-term**Asp168** | (C) electrostatic |
|  | N-term**Glu170** | (D) electrostatic, mediated by water. |
| (A) | Conv**Glu774** | (A) |
| (A), (B) | Conv**Arg721** | (A), (B) |
| (A) | Conv**Tyr722** | (B) |
| (A), (C) | Conv**Leu770** | (B) |
|  | SH1**Pro710** | (D) |
| (C) electrostatic | Conv**Asn711** | (C) electrostatic, (D) |
| (C) electrostatic, backbone (CO) | Conv**Arg712** | (C) electrostatic, backbone (CO)<br>(D) |
| (A), (B), (C) | Conv**Ile713** | (B) |
| (C) | Transducer**His666** | (D) stacking |
|  | Relay**His492** | (D) |

### Material and methods

#### Protein purification

Bovine cardiac S1 fragment was purified from fresh heart (Pel-Freez Biologicals) via the protocol described previously^4,1^. S1 was obtained by limited chymotryptic digestion of the full-length myosin in the buffer (20 mM K-Pipes, 10 mM K-EDTA, 1 mM EDTA, pH 6.8). The mix containing the protease (tosyl-Lysyl-chloromethane hydrochloride (TLCK)-treated from Sigma) was incubated at 22°C for 30 minutes. The digestion was stopped by the addition of 1 mM phenylmethylsulfonyl fluoride (PMSF). Centrifugation at 29,000 *g* during 30 minutes at 4°C allowed to remove the insoluble myosin rods. The S1 fragment was precipitated with ammonium acetate (60% w/v final) and centrifugation (29,000 x *g* during 30 minutes at 4°C). The pellet was resuspended and dialyzed with a low-salt buffer (12 mM K-Pipes, 2 mM MgCl_2_, 1 mM DTT, 0.1 mM PMSF, pH 6.8). An anion-exchange chromatography step on Mono-Q (GE Healthcare) was performed at 4°C in the buffer 20 mM Tris-HCl, 0.8 mM NaN_3_, pH 8 with a 0-350 mM NaCl gradient. Fractions containing the purified S1 were pooled and buffer-exchanged in the buffer 10 mM HEPES, 50 mM NaCl, 1 mM NaN_3_, 2.5 MgCl_2_, 0.2 mM ATP, 1 mM TCEP, pH 7.5. S1 was finally concentrated at 42 mg.ml^−1^ and 2 mM MgADP was added. The protein was aliquoted and flash frozen in liquid nitrogen for storage.

#### Mavacamten synthesis

Mava was synthesized in H.-J. Knölker’s lab (see procedure below) and also purchased from Selleck Chemicals (www.selleckchem.com). The Mava powder was dissolved in DMSO at a stock concentration of 50 mM.

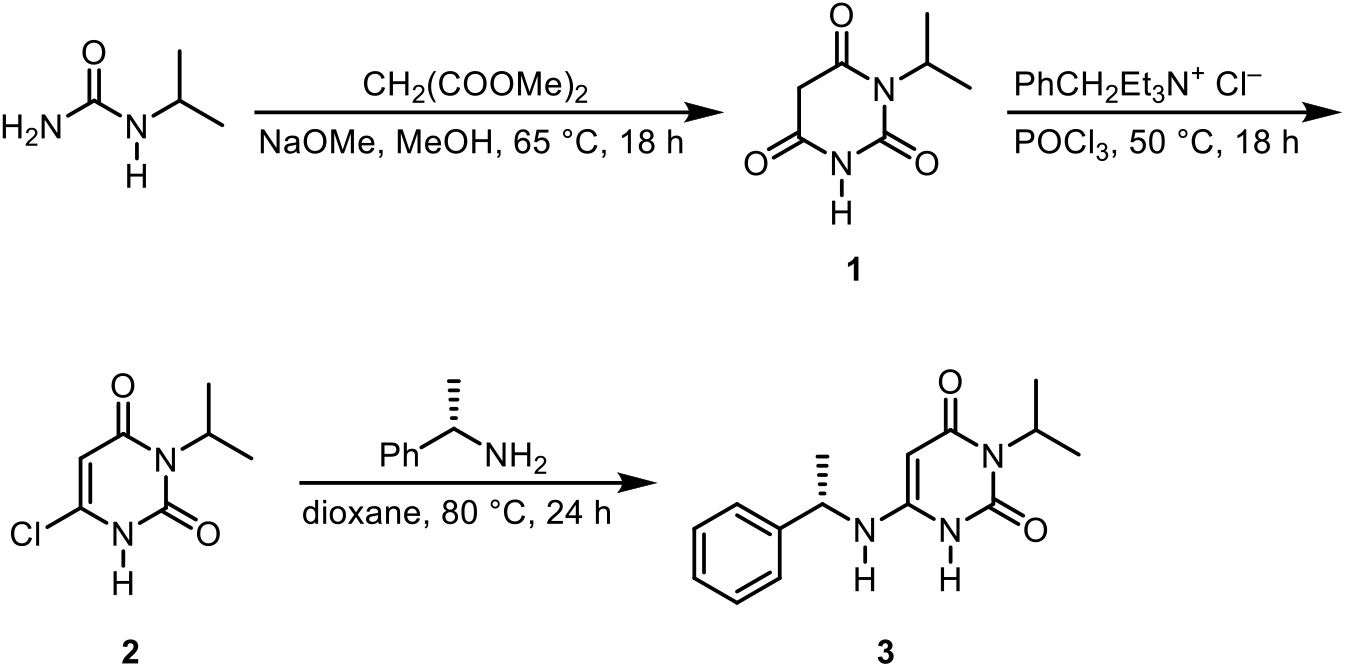

1-Isopropylbarbituric acid (1-isopropylpyrimidine-2,4,6(1*H*,3*H*,5*H*)-trione) (**1**)^5^

Dimethyl malonate (19.6 g, 148 mmol) and then sodium methanolate (18.2 g, 337 mmol) were added under continuous stirring to a solution of 1-isopropylurea (14.4 g, 141 mmol) in methanol (500 mL) at room temperature. The reaction mixture was heated under reflux for 18 h. Subsequently, the mixture was cooled first to room temperature, then to 0 °C, and acidified with hydrochloric acid (pH = 3). After removal of the solvent in vacuum, the residue was treated with ethanol (200 mL) and the resulting mixture was filtered. The filtrate was evaporated in vacuum and the residue was purified by column chomatography on silica gel (dichloromethane/methanol, 20:1) to afford 1-isopropylbarbituric acid (13.9 g, 58%) as a light yellow oil.

6-Chloro-3-isopropylpyrimidine-2,4(1*H*,3*H*)-dione (**2**) ^5^

Phosphorus oxychloride (37 mL) and benzyltriethylammonium chloride (26.1 g, 115 mmol) were added to 1-isopropylbarbituric acid (13.9 g, 81.7 mmol) under an argon atmosphere. The resulting mixture was heated at 50 °C for 18 h under an argon atmosphere. After cooling to room temperature, the excess phosphorus oxychloride was evaporated in vacuum. The residual red oil was dissolved in dichloromethane (185 mL) and water (120 mL) was added slowly over a period of 1.5 h. The layers were separated, the organic layer was washed with water (120 mL), and dried with sodium sulfate. After removal of the solvent, the residue was purified by column chomatography on silica gel (gradient elution with isohexane/ethyl acetate, from 5:1 to 1:1) to provide 6-chloro-3-isopropylpyrimidine-2,4(1*H*,3*H*)-dione (3.05 g, 20%) as a colorless solid.

(*S*)-3-Isopropyl-6-((1-phenylethyl)amino)pyrimidine-2,4(1*H*,3*H*)-dione (Mavacamten) (**3**) ^5^

(*S*)-1-Phenylethan-1-amine (1.50 mL, 1.41 g, 11.6 mmol) was added to a solution of 6-chloro-3-isopropylpyrimidine-2,4(1*H*,3*H*)-dione (1.00 g, 5.30 mmol) in 1,4-dioxane (20 mL) and the resulting solution was heated at 80 °C for 24 h under an argon atmosphere. After removal of the solvent in vacuum, the residue was dissolved in ethyl acetate (70 mL), washed with 1N HCl (2 × 50 mL), and then with a saturated aqueous solution of sodium chloride (40 mL). The organic layer was dried with sodium sulfate and the solution was concentrated in vacuum to half of the original volume in order to induce the formation of a precipitation. Hexane (20 mL) was added and the mixture was stirred at room temperature for 10 min. The resulting solid was separated by filtration, washed with hexane (20 mL), and dried in vacuum. The crude product was purified by column chomatography on silica gel (gradient elution with isohexane/ethyl acetate, from 1:1 to 1:3) to afford (*S*)-3-isopropyl-6-((1-phenylethyl)amino)pyrimidine-2,4(1*H*,3*H*)-dione (439 mg, 30%) as a colorless solid.

The purity of the compound was checked by ^1^H NMR and GC-MS.

#### Crystallization and data processing

Crystals of PPS-MD-Apo (type A) were obtained at 17°C by the siting drop vapor diffusion method from a 1:1 (v:v) ratio of protein (20 mg.ml^−1^) with 2 mM Mg.ADP.Vanadate, trypsin at a ratio 1:1000 (w:w) and precipitant containing 0.5 M Lithium sulfate, 0.1 M Tris pH 8.5 and 25% PEG 3350. Crystals of PPS-S1-Mava (type B) were obtained at 4°C by the hanging drop vapor diffusion method from a 1:1 (v:v) ratio of the protein (10 mg.ml^−1^) with Mg.ADP.BeFx, 5 mM Mavacamten, trypsin at a ratio 1:500 (w:w) and precipitant 20.5% PEG3350, 7.5% NaTacsimate pH 6.0, 5 mM TCEP. Crystals of PPS-MD-Mava (type C) were obtained at 25°C by the hanging drop vapor diffusion method from a 1:1 (v:v) ratio of the protein (10 mg.ml^−1^) with 2 mM Mg.ADP.Vanadate, 0.5 mM Mavacamten, trypsin at a ratio 1:500 (w:w) and precipitant 23% PEG3350, 0.2 M Lithium sulfate, 0.1 M Tris pH 7.9. PPS-S1-OM (type D) was reprocessed from previous data, the crystallization conditions are documented in^4^.

Crystals were transferred in the mother liquor containing 30% glycerol before and flash frozen in liquid nitrogen. X-ray diffraction data were collected at the SOLEIL synchrotron (PX2A beamline, λ = 0.98007 Å and λ = 0.9762 Å for type A and D respectively; PX1 beamline, λ = 0.98400 Å and λ = 0.97856 Å for type B and C respectively), at 100 K. Diffraction data were processed using the XDS package^6^ and AutoPROC^7^. The reprocessing of data from crystal type D was performed with the same Rfree set as the one used in 5N69. Crystals type A belongs to the P4_3_22 space group, crystals type 2 belongs to the space group P2_1_, crystals type C and D belong to the P2_1_2_1_2_1_ space group; with one molecule per asymmetric unit for type A and C and two molecules per asymmetric unit. The data collection and refinement statistics are presented in **Table 1**.

#### Structure determination and refinement

Molecular replacement was performed with bovine β-cardiac myosin (PDB code 5N69)^4^ without water and ligand as a target model with Phaser^8^ using: the motor domain (residues 80-710) for crystals type A and C and the entire S1 fragment for crystals type B and D. Manual model building was performed using Coot^9^. Drug building and restrain libraries generations were achieved using and ReadySet! from the phenix Suite^10,11^. Refinement was performed using Buster^12^ with the highest resolution structure PPS-MD-Mava as a target structure. The statistics for most favored, allowed and outlier in Ramachandran angles are for each crystal type respectively (in %): 95.64, 4.07 and 0.29 for PPS-Apo-MD; 97.58, 2.25 and 0.17 for PPS-S1-OM; 96.14, 3.30 and 0.57 for PPS-S1-Mava; 97.40, 2.45 and 0.14 for PPS-MD-Mava.

#### Molecular dynamics

Molecular dynamics procedure is similar to ^13,14^, starting from the PPS-MD-Apo, PPS-S1-Mava and PPS-S1-OM crystal structures. All molecular dynamics simulations were performed with Gromacs (version 2018.3)^15^ on all-atom systems parametrized with charmm36m forcefield^16^ and built with CHARMM-GUI^17,18^, that uses the following potential energy function (1) (see details in^19^):

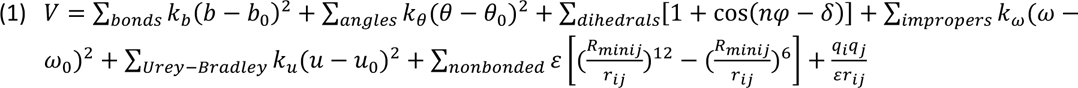

The Mg^2+^, ADP and Pi were modelled after Mg.ADP.Vanadate. All systems consisted in a box containing explicit waters TIP3 (97834 molecules), 150 mM KCl and a pH set to 7.0 with no protonation of His residues. The box consisted in a cube of 149 Å as a length of the edge and a volume of 3307949 Å^3^. Long range interactions were handled using the particle mesh Ewald (PME) method^20^ that allows to facilitation the calculation of the energies. The Coulomb sum (E(coul)) (2) is split into direct space sum (E(dir)) (3) and reciprocal space sum (E(rec)) (4) (see^21^) as follows:

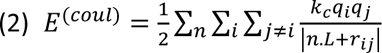

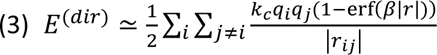

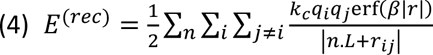

Where r_ij_ is the interparticle distance, kc is Coulomb’s constant and β is the Ewald coefficient. The electrostatic energy is considered as a periodic system expressed as a sum over all pair of interactions. While the expression of Coulomb sum is conditionally convergent, the direct and reciprocal sums are rapidly convergent sums. The Ewald formulation is used in molecular simulations to express the electrostatic energy and force (see charm-gui manual for detail, https://www.charmm-gui.org/charmmgui.org/charmmdoc/ewald.html).

The simulations were performed in an NPT ensemble. The temperature and the pressure of the system were fixed at 310.15 K with the Nosé-Hoover thermostat and 1 bar with the Parrinello-Rahman barostat^22,23^. The total duration of the simulations were 720 ns for Apo; 1020 ns for OM and Mava. The simulations were performed at least twice for each condition in order to ensure the reproducibility of the phenomenon discussed in this work. Trajectories were generated and analyzed in PyMOL^24^ which served to create the illustrations. Atomic displacements were computed with VMD^25^.

#### Frame analysis

Trajectories were assembled with ‘gmx trjconv’ from Gromacs . Each trajectory was adjusted on the first frame by superimposing the atoms of the motor backbone using macros from VMD (RMSD trajectory tool), thus allowing to extract the RMSD plots. The first images were oriented in a previous alignment so that all representations are in the same referential.

#### Binding pockets analysis

The drug binding pockets were analyzed manually. The plot representations in figures have been achieved using the software LigPlot+^26^.

#### Free energy calculations

The calculations of binding free energy (ΔGs) was performed with the software Prodigy^2,3^ at 25°C for all the frames of the time course of the simulations. The Kd was derived from the equation (5).

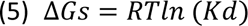

The Kd was then plotted as a function of time for the two conditions OM and Mava (**Supplementary Fig. 11**).

#### Homology modelling

Homology modelling of *O. cuniculus* SkMyo2 PPS state binding OM has been performed with SwissModel^26^ with the structure of PPS-S1-OM as a template. The figure with Mava was performed with the same model. The quality of the model has been checked by a comparison with the available structure of SkMyo2 (PDB code 6YSY ^27^).

### Supplementary Movies

**<u>Supplementary Figure 1 – Comparison of the conformation of the Lever arm in the different structures.</u>** PPS-S1-Mava is colored in cyan. PPS-S1-OM is colored in yellow. PPS-MD-Apo is colored in orange. For comparison, the last helix of the Converter from the PR state (PDB code 6FSA^1^) is shown. The specific position of Ala767 is colored in red and represented as stick.

**<u>Supplementary Movie 2 – Molecular dynamics of the PPS-S1-Apo condition (conformations explored without drug bound).</u>** Different views are presented: the overall view to appreciate the dynamics of the Lever arm; the drug binding pocket (drug targeted site) in two orientations; the catalytic site. The subdomains are colored differently: N-terminal extension is colored in pink; N-terminal subdomain in grey; U50 in dark blue; L50 in light orange; Relay in yellow; SH1-helix in red; Converter in green; IQ region in cyan; ELC in light pink. Switch-1 (bright pink), Switch-2 (orange) are two connectors close to the catalytic site.

**<u>Supplementary Movie 3 – Molecular dynamics of the PPS-S1-OM (conformations explored when OM is bound).</u>** Different views are presented: the overall view to appreciate the dynamics of the Lever arm; the drug binding pocket (drug targeted site) in two orientations; the catalytic site. The subdomains are colored differently: N-terminal extension is colored in pink; N-terminal subdomain in grey; U50 in dark blue; L50 in light orange; Relay in yellow; SH1-helix in red; Converter in green; IQ region in cyan; ELC in light pink. Switch-1 (bright pink), Switch-2 (orange) are two connectors close to the catalytic site.

**<u>Supplementary Movie 4 – Molecular dynamics of the PPS-S1-Mava (conformations explored when Mava is bound)</u>**. Different views are presented: the overall view to appreciate the dynamics of the Lever arm; the drug binding pocket (drug targeted site) in two orientations; the catalytic site. The subdomains are colored differently: N-terminal extension is colored in pink; N-terminal subdomain in grey; U50 in dark blue; L50 in light orange; Relay in yellow; SH1-helix in red; Converter in green; IQ region in cyan; ELC in light pink. Switch-1 (bright pink), Switch-2 (orange) are two connectors close to the catalytic site.

**<u>Supplementary Movie 5 – Dynamics of OM in the drug binding pocket during the time course of the molecular dynamics simulation.</u>** The movie compares the initial position (structure, orange) of PPS-OM-S1 to the evolution during the time course of the dynamics (green). The nucleotide (yellow) and OM (dark blue) are represented as spheres.

**<u>Supplementary Movie 6 – Dynamics of Mava in the drug binding pocket during the time course of the molecular dynamics simulation.</u>** The movie compares the initial position (structure, orange) of PPS-Mava-S1 to the evolution during the time course of the dynamics (green). The nucleotide (yellow) and Mava are represented as spheres. Mava is colored in dark blue in the initial position (structure) and in yellow in the dynamics.

**<u>Supplementary Movie 7 – Dynamics of OM and Mava in the drug binding site.</u>** The movie compares how OM and Mava explore the drug binding pocket during the time course of the dynamics. The two movies were synchronized to compare the positions at the same time.

